# Stereoselective Covalent Inhibitor of the Ovarian Cancer-Driving Transcription Factor PAX8

**DOI:** 10.64898/2026.08.15.745031

**Authors:** Taylor M. Nuttall, Aman Modi, Kelvin Li, Emily A. Lau, Alicia Zhang, Bhavna Malik, Tezcan Guney, John Eksterowicz, Gregory T. Notte, Thomas J. Maimone, Daniel K. Nomura

**Affiliations:** Departments of Chemistry and Molecular and Cell Biology, University of California, Berkeley, Berkeley, CA 94720 USA; Innovative Genomics Institute, Berkeley, CA 94720 USA; Molecular Therapeutics Initiative, Berkeley, CA 94720 USA; Gilead Sciences Inc., Foster City, CA 94404 USA

**Keywords:** PAX8, ovarian cancer, chemoproteomics, activity-based protein profiling, cysteine, covalent drug discovery

## Abstract

Transcription factors remain among the most challenging therapeutic targets in part because they lack well-defined ligandable binding pockets. We recently showed that aberrantly reactive cysteines in transcription factors can be directly targeted with electrophilic small molecules to induce selective transcription factor destabilization and degradation. Here, we extend this strategy to the lineage-defining oncogenic transcription factor PAX8, a critical driver of ovarian cancer. Screening of a library of cysteine-reactive compounds against an endogenously HiBiT-tagged PAX8 reporter identified a sulfinyl aziridine chemotype that selectively reduced PAX8 abundance. Structure-activity and stereochemical analyses revealed highly enantio- and diastereoselective activity, identifying KL6-159A as the lead compound. Quantitative proteomics demonstrated selective loss of PAX8, while cellular thermal shift analysis and chemoproteomic profiling established direct covalent engagement of PAX8 at cysteine C57. Mutation of C57 completely abolished KL6-159A-induced PAX8 depletion, demonstrating that this residue is essential for compound activity. Transcriptomic profiling revealed broad suppression of the PAX8 transcriptional program, with FOXM1 emerging as the most significantly downregulated regulatory network together with numerous established PAX8 target genes. Collectively, these studies establish direct covalent engagement, transcriptional inhibition, and destabilization of PAX8 and further demonstrate the generality of covalent chemoproteomic approaches for drugging previously intractable transcription factors.

## Introduction

Transcription factors are among the largest remaining classes of therapeutically important but poorly druggable proteins in modern-day drug discovery. Despite their central roles in development, lineage specification, and human disease, most transcription factors have resisted conventional small-molecule drug discovery because they generally lack well-defined ligand-binding pockets and instead rely heavily on large protein-protein and protein-DNA interaction interfaces, many of which are embedded within intrinsically disordered regions (IDRs) ^1,2^. Consequently, transcription factors have historically been considered intractable to conventional pharmacological targeting, limiting therapeutic intervention against many of the oncogenic drivers that sustain human cancers. While recent advances in targeted protein degradation have expanded the range of proteins amenable to pharmacological modulation, including against transcription factors, many degrader strategies remain dependent upon first identifying a ligand capable of directly engaging the target protein ^3,4^. Thus, discovering methods to directly target previously inaccessible transcription factors remains a major challenge in chemical biology.

Recent studies have started to demonstrate that even intrinsically disordered transcription factors are not necessarily devoid of chemically tractable binding sites. Rather, dynamic conformational ensembles within IDRs can harbor ligandable amino acid residues that enable selective small-molecule engagement. We and others have shown that reactive cysteine residues embedded within intrinsically disordered proteins can serve as privileged sites for covalent ligand discovery using chemoproteomic approaches ^5–11^. Using this strategy, our laboratory has directly targeted several previously undruggable transcription factors, including MYC, CTNNB1, AR/AR-V7, and IRF5/IRF8, through covalent engagement of intrinsically disordered cysteines, resulting in selective destabilization and degradation of these proteins ^7–11^. These findings suggest that exploiting reactive cysteines is a broadly applicable strategy to directly drug transcription factors that have historically been inaccessible to small-molecule modulation.

Among these challenging targets, the paired box transcription factor PAX8 represents a particularly compelling therapeutic opportunity. PAX8 is essential for embryonic development of the thyroid gland, kidney, and Müllerian tract, where it controls lineage specification and tissue differentiation. While PAX8 expression is largely restricted in adult tissues, aberrant reactivation or sustained PAX8 expression is a hallmark of multiple malignancies ^12–16^. Most notably, PAX8 functions as a lineage-survival transcription factor in high-grade serous ovarian carcinoma (HGSOC), where genetic depletion of PAX8 profoundly inhibits tumor cell proliferation and survival. Beyond ovarian cancer, PAX8 has also been implicated in fallopian tube carcinoma, primary peritoneal carcinoma, ovarian endometrioid carcinoma, thyroid carcinoma, renal cell carcinoma, thymic epithelial tumors, and subsets of endometrial and pancreatic cancers. Mechanistically, PAX8 regulates extensive transcriptional programs controlling cell-cycle progression, chromatin organization, metabolism, lineage identity, and metastatic behavior, in part by maintaining downstream oncogenic transcriptional regulators, including FOXM1 ^12–17^. Consequently, PAX8 has emerged as one of the most attractive lineage-specific therapeutic targets in ovarian cancer. However, despite considerable biological validation, no direct small-molecule inhibitors or degraders of PAX8 have been publicly reported.

Here, we used covalent chemoproteomic approaches to directly discover small molecules that engage and destabilize PAX8. Screening a chemically diverse electrophile library comprising over 3,000 cysteine-reactive molecules, including acrylamides, chloroacetamides, and small strained rings identified a chiral sulfinyl aziridine that reduced endogenous PAX8 abundance in manner dependent on compound stereochemistry. Through complementary chemoproteomic, biochemical, and genetic studies, we demonstrate that the lead compound KL6-159A directly and covalently engages C57 within PAX8, resulting in selective protein destabilization and suppression of PAX8 transcriptional activity. Transcriptomic profiling further revealed modulation of the PAX8-regulated transcriptional network, with FOXM1 emerging as the most significantly affected downstream regulatory program. Collectively, these studies establish direct covalent destabilization of PAX8 through targeting a reactive cysteine as a tractable strategy for targeting this important oncogenic transcription factor.

## Results

### Covalent Chemoproteomic Screening Identifies a Stereoselective Covalent Destabilizer of PAX8

Given our previous success in directly targeting transcription factors cysteines ^7–11^, we sought to determine whether a similar strategy could be applied to PAX8. To identify small molecules capable of directly destabilizing PAX8, we screened a chemically diverse library of more than 3,000 cysteine-reactive electrophiles, including acrylamides, chloroacetamides, and sulfinyl aziridines,^8^ using an OVCAR3 ovarian cancer cell line expressing an endogenous N-terminal HiBiT-tagged PAX8 reporter. Three compounds, KL6-159A, KL6-167A, and KL6-172A, reduced HiBiT-PAX8 abundance by greater than 70%, identifying a common sulfinyl aziridine oxindole scaffold as a promising chemotype for PAX8 modulation **(Figure 1a)**.

**Figure 1.**
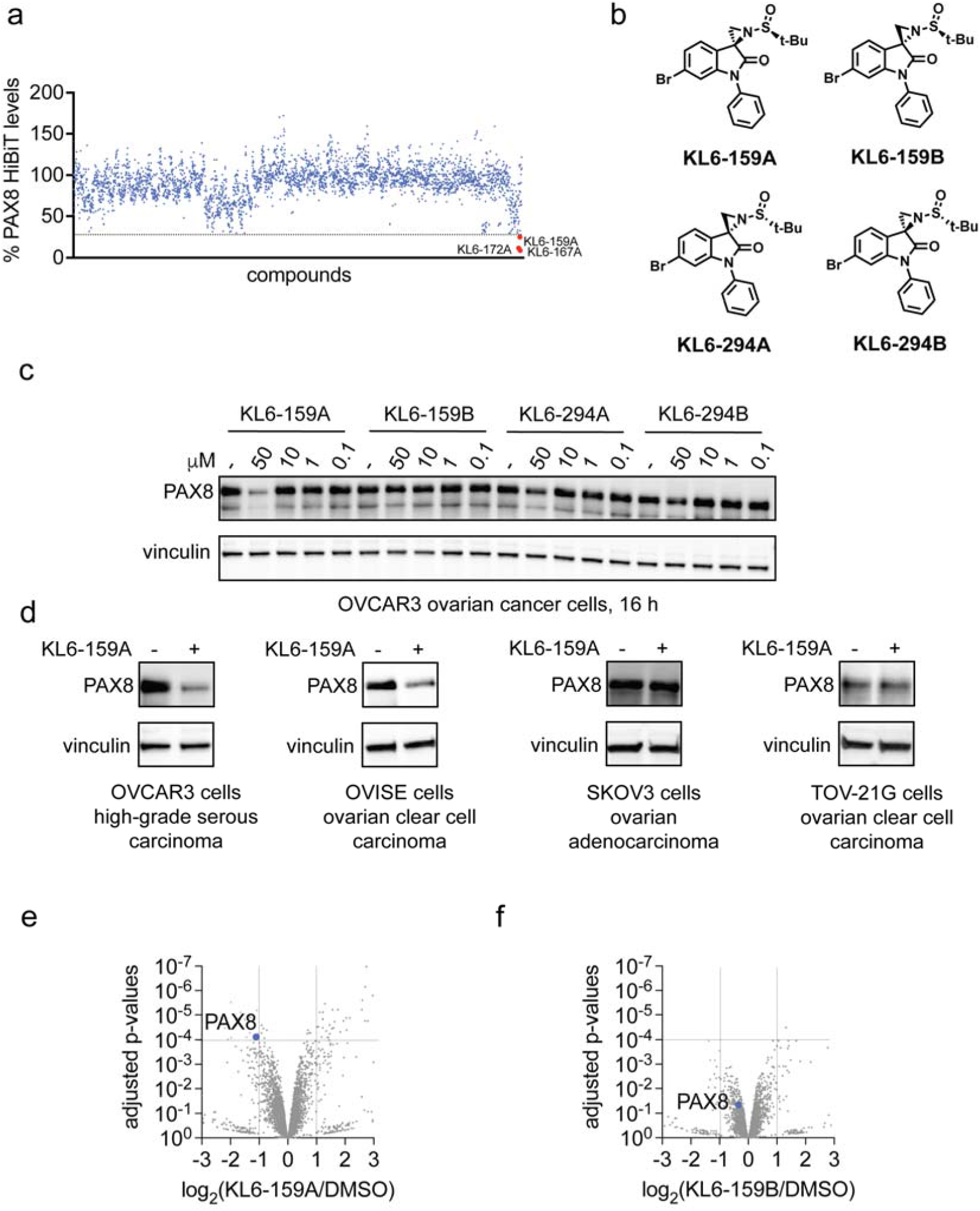
Covalent chemoproteomic screening identifies a stereoselective covalent destabilizer of PAX8. **(a)** Screening of a library of >3,000 cysteine-reactive electrophiles in OVCAR3 ovarian cancer cells expressing endogenously N-terminal HiBiT-tagged PAX8 identifies three sulfinyl aziridine hits (KL6-172A, KL6-159A, and KL6-167A) that reduce PAX8 abundance by >70%. **(b)** Chemical structures of KL6-159A, its diastereomer KL6-159B, and the corresponding stereoisomers KL6-294A and KL6-294B. **(c)** Endogenous PAX8 levels in OVCAR3 cells treated with DMSO vehicle or indicated compounds for 16 h, as assessed by SDS/PAGE and Western blotting. Vinculin serves as a loading control. **(d)** KL6-159A reduces endogenous PAX8 levels in OVCAR3 and OVISE cells but not in SKOV3 or TOV-21G cells. Cells were treated with DMSO vehicle or KL6-159A (50 µM, 16 h for OVCAR3, OVISE, and SKOV3 cells and 24 h for TOV-21G cells), after which PAX8 and the loading control vinculin levels were assessed by SDS/PAGE and Western blotting. **(e**,**f)** Tandem mass tag (TMT)-based quantitative proteomic profiling demonstrates selective reduction of PAX8 following treatment with KL6-159A **(e)**, whereas the inactive enantiomer KL6-159B does not significantly alter PAX8 abundance **(f)**. OVISE cells were treated with KL6-159A or KL6-159B (50 µM, 16 h), after which proteomes were analyzed by LC-MS/MS. Volcano plots depict log_2_ protein abundance changes relative to vehicle-treated controls. Screening data for **(a)** can be found in **Table S1**. Blots in **(c**,**d)** are representative of n=3 biologically independent replicates per group. Proteomics data in **(e**,**f)** are in **Table S2** and come from n=3 biologically independent replicates per group.

Because stereoselective engagement suggests specific molecular recognition rather than nonspecific electrophile reactivity, we next evaluated each active compound along with its corresponding diastereomer. All three stereochemical pairs exhibited stereoselective reduction of HiBiT-PAX8, with KL6-159A displaying the largest difference in activity relative to its diastereomer KL6-159B **(Figure S1)**. Encouraged by these findings, we further examined the complete stereochemical series consisting of the diastereomeric pair KL6-159A/KL6-159B and the corresponding diastereomeric pair KL6-294A/KL6-294B **(Figure 1b)**. Remarkably, only KL6-159A reduced endogenous PAX8 protein abundance in OVCAR3 cells, whereas its stereoisomers were essentially inactive, demonstrating a high degree of stereochemical specificity **(Figure 1c)**. These data strongly suggested that KL6-159A acts through a highly specific target engagement mechanism rather than nonspecific protein modification.

We next evaluated whether KL6-159A exhibited activity across multiple ovarian cancer models. KL6-159A substantially reduced endogenous PAX8 protein levels in the high-grade serous ovarian carcinoma cell line OVCAR3 and the ovarian clear cell carcinoma cell line OVISE but failed to reduce PAX8 abundance in SKOV3 ovarian adenocarcinoma cells or TOV-21G ovarian clear cell carcinoma cells **(Figure 1d)**. Thus, KL6-159A displays subtype-selective activity among ovarian cancer models, suggesting that additional cellular determinants beyond PAX8 expression may influence KL6-159A-induced PAX8 depletion. These results may also suggest that while KL6-159A may still engage PAX8 in these cells, the machinery responsible for PAX8 loss may not be found in these cell lines.

To determine the overall proteome-wide selectivity of KL6-159A-mediated PAX8 loss, we performed tandem mass tag (TMT)-based quantitative proteomic profiling following treatment of OVISE cells with KL6-159A or its inactive diastereomer KL6-159B. KL6-159A selectively reduced PAX8 abundance while only 15 additional proteins were significantly downregulated across the quantified proteome **(Figure 1e)**. In contrast, KL6-159B did not significantly reduce PAX8 levels **(Figure 1f)**, showing that proteome-wide activity is likewise stereoselective. Together, these studies identify KL6-159A as a highly stereoselective and proteome-selective covalent modulator of PAX8.

### KL6-159A Directly and Covalently Engages PAX8 Through C57 to Induce Protein Destabilization

Having identified a molecule that selectively induces PAX8 depletion, we next sought to determine whether the compound directly engaged PAX8. Because our previous covalent destabilizing ligands for MYC, CTNNB1, AR/AR-V7, IRF5, and IRF8 directly destabilized their target proteins upon covalent engagement ^7–11^, we first assessed the effects of KL6-159A on PAX8 thermal stability using cellular thermal shift analysis (CETSA) ^18^. KL6-159A markedly destabilized PAX8 in OVCAR3 cells, decreasing the apparent melting temperature of PAX8 from 57.5 °C to 51.1 °C, consistent with direct compound binding and ligand-induced destabilization **(Figure 2a-2b)**.

**Figure 2.**
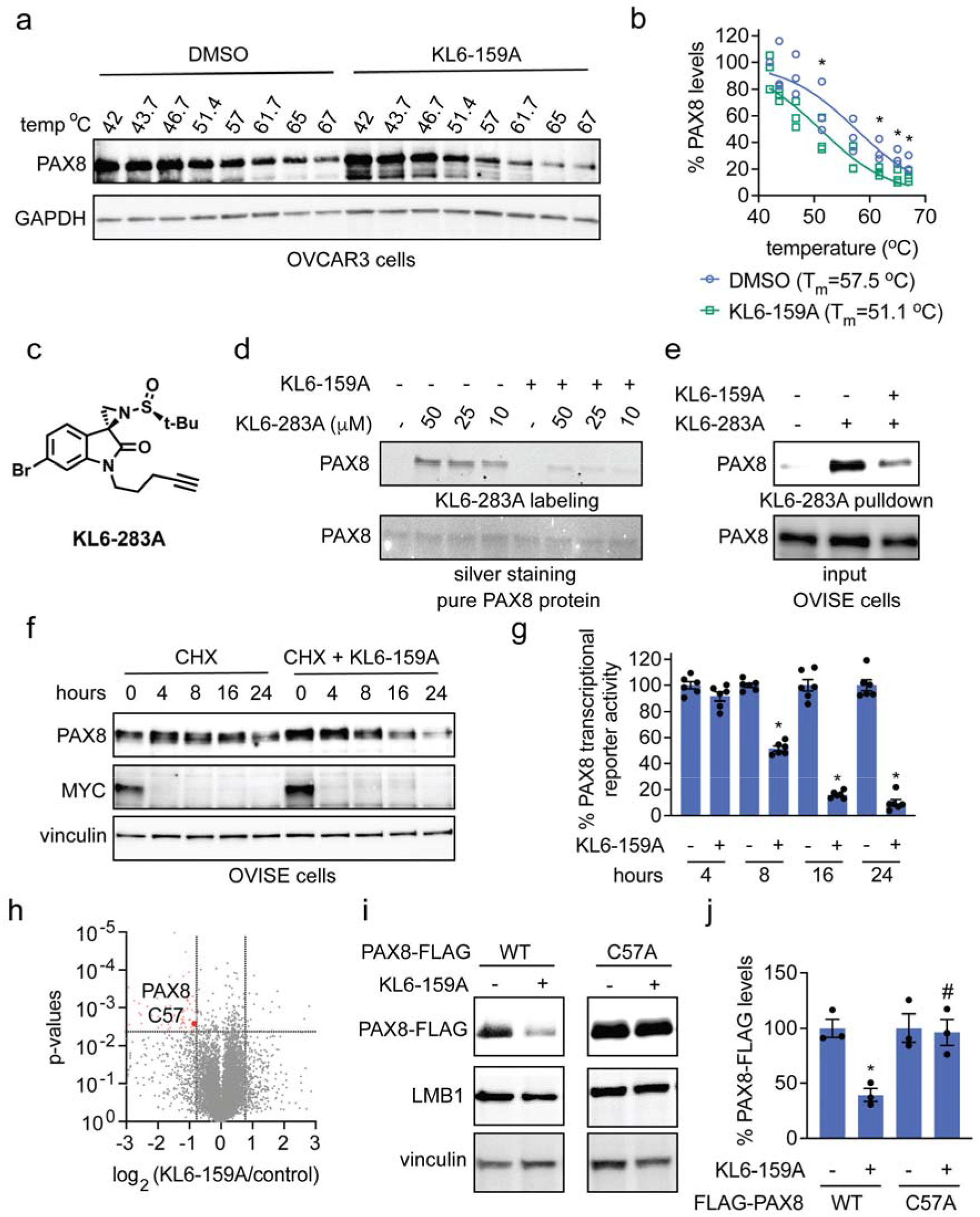
KL6-159A directly engages PAX8 through C57 to induce protein destabilization and inhibition of transcriptional activity. **(a,b)** CETSA demonstrating ligand-induced destabilization of endogenous PAX8 following DMSO vehicle or KL6-159A (50 µM, 4 h) treatment in OVCAR3 cells. We centrifuged the resulting lysates to clear aggregated proteins, then assessed PAX8 and GAPDH control levels by SDS/PAGE and Western blotting. **(c)** Structure of the alkyne-functionalized probe KL6-283A derived from KL6-159A. **(d)** Dose-dependent covalent labeling of purified recombinant PAX8 by KL6-283A that is competed by excess KL6-159A. PAX8 protein (0.5 µg) was pre-incubated with DMSO vehicle or KL6-159A (150 µM) for 30 min at 37 °C prior to labeling with the indicated concentrations of KL6-283A for 30 min, after which probe-labeled proteins were subjected to copper-catalyzed azide-alkyne cycloaddition (CuAAC)-mediated appendage of azide-functionalized rhodamine, after which probe labeling was assessed by SDS/PAGE and fluorescence visualization, with silver staining to assess loading. **(e)** KL6-283A enriches endogenous PAX8 from OVISE cells, and enrichment is partially outcompeted by pretreatment with KL6-159A. OVISE cells were pre-incubated with DMSO vehicle or KL6-159A (100 µM) for 30 min prior to labeling with KL6-283A (25 µM) for 1 h, after which probe-labeled proteins were subjected to CuAAC-mediated appendage of an azide-functionalized biotin enrichment handle, avidin-enrichment, elution, and assessment by SDS/PAGE and Western blotting for PAX8. An aliquot of the pre-enrichment sample was run to assess equivalent input. **(f)** Cycloheximide (CHX) chase experiments indicate that KL6-159A accelerates PAX8 loss without inhibiting protein synthesis. MYC is shown as a short-lived protein control. OVISE cells were pre-treated with DMSO vehicle or cycloheximide (100 µg/mL) for 1 h prior to treatment of cells with DMSO vehicle or KL6-159A (50 µM) for the indicated time, and PAX8, MYC, and vinculin levels were assessed by SDS/PAGE and Western blotting. **(g)** KL6-159A time-dependently suppresses PAX8-dependent luciferase reporter activity. OVISE cells transduced with a PAX8 luciferase reporter construct by lentivirus were treated with DMSO or KL6-159A (50 µM), and PAX8 transcriptional reporter activity was assessed by luminescence. **(h)** isoDTB-ABPP identifies selective covalent engagement of PAX8 C57 with limited proteome-wide off-target engagement. OVISE cells were treated with DMSO vehicle or KL6-159A (50 µM) for 2 h, after which the resulting lysates were labeled with a cysteine-reactive alkyne-functionalized iodoacetamide probe and taken through the isoDTB-ABPP method. **(i**,**j)** Mutation of C57 to alanine abolishes KL6-159A-induced loss of nuclear PAX8-FLAG in HEK293T cells. HEK293T cells expressing PAX8-FLAG were treated with DMSO vehicle or KL6-159A (50 µM) for 4 h, after which nuclear fractions were isolated, and PAX8-FLAG, the loading control LMB1, and vinculin were assessed by SDS/PAGE and Western blotting. Blots in **(a**,**d**,**e**,**f**,**i)** are representative of n=3 biologically independent replicates per group. **(b)** shows individual replicate values from the experiment described in **(a). (g**,**j)** show individual replicate values and average ± sem. Significance in **(b**,**g)** is expressed as *p<0.05 compared to vehicle-treated controls at each temperature or time point, respectively. Significance in **(j)** is expressed as *p<0.05 compared to respective vehicle-treated controls and #p<0.05 compared to KL6-159A-treated WT controls. Data in **(h)** are in **Table S3**.

To directly assess covalent engagement of PAX8, we synthesized an alkyne-functionalized analog of KL6-159A, termed KL6-283A, suitable for copper-catalyzed azide-alkyne cycloaddition click chemistry-based detection **(Figure 2c)**. KL6-283A labeled purified recombinant PAX8 protein in a concentration-dependent manner, and labeling was effectively outcompeted by excess KL6-159A, demonstrating direct covalent engagement of PAX8 by the parent compound **(Figure 2d)**. Consistent with these observations, KL6-283A also enriched endogenous PAX8 from OVISE cells, and pretreatment with KL6-159A partially outcompeted this enrichment, confirming target engagement in cells **(Figure 2e)**. These studies establish that KL6-159A directly binds PAX8 both *in vitro* and in intact cells.

To determine whether KL6-159A reduced PAX8 protein levels by inhibiting protein synthesis, we performed cycloheximide chase experiments in OVISE cells. Treatment with cycloheximide alone resulted in gradual loss of PAX8 over time, whereas co-treatment with KL6-159A accelerated the disappearance of PAX8 protein without affecting translational inhibition, indicating that KL6-159A promotes loss of PAX8 at the protein level rather than by suppressing protein synthesis **(Figure 2f)**. Consistent with reduced PAX8 abundance, KL6-159A also produced a progressive inhibition of PAX8-dependent transcriptional reporter activity over 24 hours, demonstrating that compound-mediated destabilization results in functional suppression of PAX8 transcriptional activity **(Figure 2g)**.

To identify the site of covalent modification, we next performed isotopic desthiobiotin activity-based protein profiling (isoDTB-ABPP) ^6,19,20^. Across more than 13,700 quantified cysteines, KL6-159A exhibited high proteome-wide selectivity, significantly engaging only 67 cysteines, with C57 of PAX8 emerging as a significantly occupied site **(Figure 2h)**. To determine whether this residue was required for compound activity, we generated a C57A mutant of FLAG-tagged PAX8. Whereas KL6-159A efficiently reduced wild-type nuclear PAX8-FLAG abundance, mutation of C57 completely abolished compound-induced protein loss **(Figure 2i-2j)**, demonstrating that covalent engagement of C57 is required for KL6-159A-mediated PAX8 destabilization. Collectively, these data establish that KL6-159A directly and covalently engages PAX8 through C57 to destabilize and degrade the protein.

### Direct Covalent Destabilization of PAX8 Suppresses the FOXM1 Transcriptional Program

To determine the downstream transcriptional consequences of direct PAX8 destabilization, we performed RNA sequencing on OVISE cells following KL6-159A treatment. KL6-159A induced widespread transcriptional remodeling, altering the expression of thousands of transcripts. Among the most significantly downregulated genes were numerous established PAX8-regulated genes, including WNT7A, FGF18, FGFR2, CCND1, ITGB3, CCNB1, CCNB2, CDK2, SOX2, and CXCL8^17,21–24^ **(Figure 3a)**, consistent with broad suppression of the PAX8 transcriptional program.

**Figure 3.**
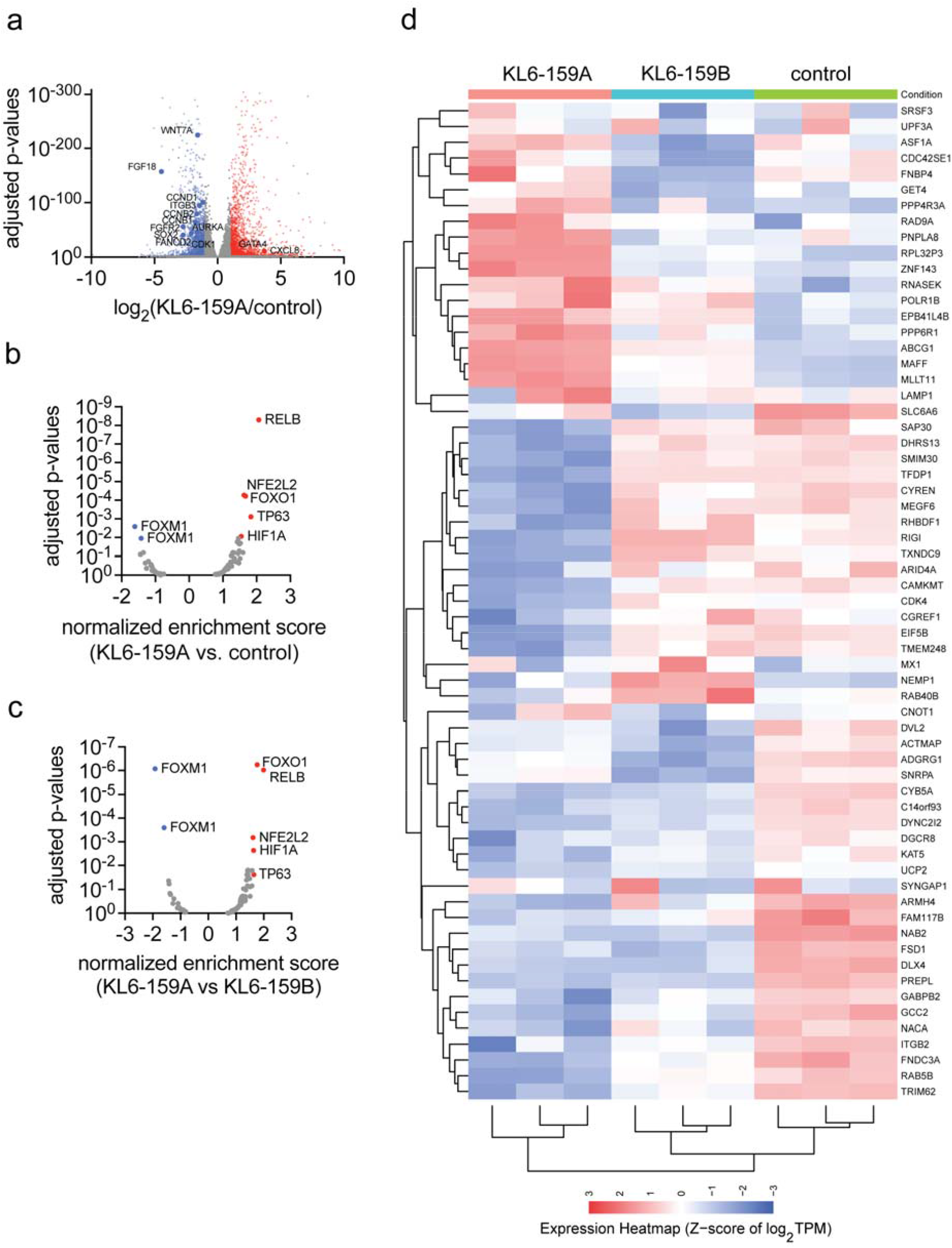
Direct covalent destabilization of PAX8 suppresses the FOXM1 transcriptional program. **(a)** RNA-sequencing analysis of OVISE ovarian cancer cells treated with KL6-159A. OVISE cells were treated with DMSO vehicle, KL6-159A, or KL6-159B (50 µM) for 16 h. **(b)** Transcription factor enrichment analysis identifies FOXM1 as the most significantly downregulated transcription factor in the network following KL6-159A treatment relative to vehicle-treated controls. **(c)** Comparison of KL6-159A with the inactive enantiomer KL6-159B demonstrates stereoselective suppression of the FOXM1 transcriptional program. **(d)** Heatmap showing representative differentially expressed genes uniquely regulated by KL6-159A compared with KL6-159B and vehicle-treated cells, illustrating stereoselective suppression of the PAX8 transcriptional program. RNA-seq data are provided in **Table S4**.

To gain insight into the transcriptional networks perturbed by KL6-159A, we performed transcription factor enrichment analysis. Strikingly, the FOXM1 transcriptional network emerged as the most significantly downregulated pathway following KL6-159A treatment compared with vehicle control **(Figure 3b)**. Moreover, comparing KL6-159A with its inactive enantiomer KL6-159B also identified FOXM1 as the most significantly suppressed transcriptional program **(Figure 3c)**, demonstrating that inhibition of FOXM1 signaling is stereoselective and directly associated with active PAX8 targeting. FOXM1 is a critical downstream driver of PAX8 transcriptional programming; these data therefore demonstrate on-target transcriptional modulation in ovarian cancer cells ^14,25^. Additional transcription factor programs, including HIF1A, TP63, FOXO1, NFE2L2, and RELB, were also significantly altered but to a lesser extent than FOXM1, suggesting that FOXM1 represents the dominant downstream transcriptional consequence of PAX8 destabilization.

Consistent with these analyses, hierarchical clustering demonstrated a distinct transcriptional profile uniquely associated with KL6-159A that was absent following treatment with the inactive enantiomer KL6-159B **(Figure 3d)**. Together, these findings show that direct covalent destabilization of PAX8 selectively collapses the PAX8 transcriptional program and identify suppression of FOXM1 signaling as a major downstream consequence of pharmacological PAX8 targeting.

## Discussion

Transcription factors have long been among the most important yet least tractable classes of therapeutic targets. Consequently, developing approaches that enable direct small-molecule targeting of previously inaccessible transcription factors remains a major objective in chemical biology. In the present study, we use a covalent chemoproteomic strategy to show that the lineage-defining oncogenic transcription factor PAX8 can be directly targeted by exploiting the reactive cysteine C57 within a folded domain of PAX8. Screening a chemically diverse library of electrophilic compounds identified a sulfinyl aziridine oxindole scaffold that selectively destabilizes PAX8 in ovarian cancer cells. We previously disclosed this novel cysteine-reactive sulfinyl aziridine warhead in a paper in which we discovered a covalent destabilizing degrader of another oncogenic transcription factor, MYC ^8^. Multiple orthogonal lines of evidence support a direct mechanism of action, including stereoselective activity, ligand-induced thermal destabilization, direct covalent engagement of purified and cellular PAX8, chemoproteomic identification of C57 as the principal site of engagement, and complete loss of activity upon mutation of C57. Together, these data establish that KL6-159A directly engages PAX8 by covalently modifying C57 to induce selective protein destabilization.

A notable feature of the direct engagement of PAX8 by KL6-159A is its exceptional stereochemical specificity. The active compound exhibited pronounced stereoselectivity relative to its stereoisomers KL6-159B, KL6-294A, and KL6-294B, which were largely inactive. This degree of stereochemical discrimination is difficult to reconcile with nonspecific electrophilic protein modification and instead indicates that productive covalent bond formation follows a highly defined molecular recognition event. Similar stereochemical behavior has emerged as a hallmark of selective covalent ligands identified through chemoproteomic approaches ^5,26^, and provides compelling evidence that KL6-159A engages a well-defined binding pose before covalent modification of C57.

Cysteine 57, alongside C45, on PAX8 has been characterized as a crucial switch for DNA binding and is highly conserved across the PAX family. Along with C45, C57 regulates the ability of PAX8 to bind to specific promoter sequences, and both of these cysteines have been found to be redox-regulated cysteines ^27^. Previous studies have shown that PAX8 loses its ability to bind DNA upon oxidation of these two cysteines ^27^. Both C45 and C57 have also been found to be ligandable in a comprehensive cysteine chemoproteomic ligandability mapping endeavor across cancer cell lines ^28^. C57 has also been shown to have a reduced pK_a_ of 7.6, indicating its hyper-reactivity ^29^. Interestingly, BridGene announced that they also developed a covalent inhibitor against PAX8 that targets C45, also within the PAX8 DNA-binding domain. The structures of their molecules, however, have not yet been published ^30^.

Our findings further expand an emerging framework in which aberrantly reactive cysteines within transcription factors serve as privileged ligandable hotspots for direct pharmacological modulation of these challenging targets. Using related covalent chemoproteomic strategies, we previously demonstrated direct destabilization and degradation of MYC, CTNNB1, AR/AR-V7, and IRF5/IRF8 by targeting reactive cysteines within these transcription factors ^7–10^. For MYC, we also demonstrated stereoselective, covalent engagement, destabilization, and degradation induced by a sulfinyl aziridine warhead ^8^. Erb and Cravatt also published on the stereoselective covalent modification of a cysteine within the transcription factor FOXA1 to rewire its transcriptional programming ^5^. The present study extends this concept to PAX8, a lineage-specifying paired-box transcription factor that has remained refractory to conventional drug discovery efforts. Collectively, these studies suggest that covalent targeting of certain reactive nucleophilic sites may represent a vulnerability within transcription factors that can be broadly exploited for pharmacological modulation and potential therapeutic benefit.

Beyond establishing direct target engagement, our transcriptomic analyses provide insight into the biological consequences of pharmacological PAX8 loss. Among all transcriptional programs altered following KL6-159A treatment, the FOXM1 regulatory network emerged as the most significantly suppressed pathway. FOXM1 is a well-established driver of cell-cycle progression, mitotic fidelity, DNA replication, and DNA repair and has previously been implicated as a critical downstream effector of PAX8 in ovarian cancer ^14,15,25^. Consistent with this model, KL6-159A coordinately downregulated numerous canonical FOXM1 target genes, along with other established PAX8-regulated genes. These findings suggest that pharmacological destabilization of PAX8 suppresses a lineage-specific transcriptional program that converges on FOXM1-mediated control of proliferation and survival. Future studies examining the mechanistic relationship between PAX8 occupancy and FOXM1 transcriptional regulation will further clarify how disruption of this regulatory axis contributes to the antitumor activity of PAX8-targeting compounds.

Although our data strongly support direct covalent engagement of PAX8 and accelerated protein destabilization, the precise degradation pathway responsible for PAX8 turnover remains to be established. Rescue experiments using proteasome, Cullin neddylation, and autophagy inhibitors were precluded by the pronounced toxicity of these agents in ovarian cancer cells, preventing definitive assignment of the degradation machinery. Nevertheless, combined CETSA, cycloheximide chase, chemoproteomic, and mutagenesis studies collectively demonstrate that KL6-159A acts through direct covalent engagement of PAX8 to promote protein destabilization rather than inhibit protein synthesis. Elucidating the degradation machinery responsible for PAX8 turnover following C57 engagement is an important direction for future investigation.

Furthermore, our current molecule lacks sufficient potency and overall selectivity and likely does not have favorable pharmacokinetic properties for *in vivo* studies. Further medicinal chemistry efforts will be required to improve potency, selectivity, and drug-like properties to enable testing of PAX8 inhibitors and degraders in pre-clinical cancer models. Finally, these studies further illustrate the power of integrating phenotypic screening with covalent chemoproteomics for ligand discovery against historically intractable targets.

In summary, we report the discovery of a stereoselective covalent small molecule that directly engages C57 within PAX8 to induce selective protein destabilization and suppression of oncogenic PAX8-dependent transcriptional programs. These findings establish PAX8 as a directly druggable transcription factor and further support the concept that cysteine-targeting represents exploitable molecular entry points for targeting previously undruggable transcription factors. More broadly, this work expands the growing repertoire of transcription factors that can be directly modulated through covalent chemoproteomic approaches and provides a general framework for discovering therapeutics against lineage-defining transcriptional dependencies in cancer.

## Acknowledgment

We thank the members of the Nomura Research Group and Gilead for critically reading the manuscript. This work was also supported by Gilead Science Inc., the National Science Foundation Molecular Foundations for Biotechnology (MFB) grant (2127788), the UC Berkeley Molecular Therapeutics Initiative (MTI), the Mark Foundation for Cancer Research ASPIRE Award, the National Institutes of Health (R35CA263814, R01CA240981), and the Bakar Award. We also thank Hasan Celik, Alicia Lund, and the UC Berkeley NMR facility in the College of Chemistry (CoC-NMR) for spectroscopic assistance. Instruments in the College of Chemistry NMR facility are partly supported by NIH S10OD024998.

## Author Contributions

TMN, GN, DKN conceived of and co-directed the project. TMN, AM, DKN performed experiments for the paper. TMN, AM, BM, TG, JE, GTN, DKN provided critical insights into project experiments. TMN, DKN wrote the paper.

