## Supporting Information for "Stereoselective Covalent Inhibitor of the Ovarian Cancer-Driving Transcription Factor PAX8"

**Supporting Table Legends**

**Table S1. Screening of cysteine-reactive covalent ligands to identify PAX8 degraders.** Screening of a library of >3,000 cysteine-reactive electrophiles in OVCAR3 ovarian cancer cells expressing endogenously N-terminal HiBiT-tagged PAX8. OVCAR3 cells were treated with DMSO vehicle or compound (25 μM, 24 h) and luminescence was read out. Shown in the table are the names and structures of all compounds tested and their activity in the screen.

**Table S2. Proteomic profiling of KL6-159A and KL6-159B in OVISE cells.** Tandem mass tag (TMT)-based quantitative proteomic profiling of OVISE cells treated with DMSO vehicle, KL6-159A (50 μM), or KL6-159B (50 μM) for 16 h, after which proteomes were analyzed by LC-MS/MS. Data are from n=3 biologically independent replicates per group.

**Table S3. Chemoproteomic profiling of KL6-159A in OVISE cells.** isoDTB-ABPP identifies selective covalent engagement of PAX8 C57 with limited proteome-wide off-target engagement. OVISE cells were treated with DMSO vehicle or KL6-159A (50 μM) for 2 h, after which the resulting lysates were labeled with a cysteine-reactive alkyne-functionalized iodoacetamide probe and taken through the isoDTB-ABPP method. Data are from n=3 replicates per group.

**Table S4. RNA-seq of KL6-159A and KL6-159B in OVISE cells.** RNA-sequencing analysis of OVISE ovarian cancer cells treated with KL6-159A. OVISE cells were treated with DMSO vehicle or KL6-159A (50 μM) for 16 h. The table contains the RNA-seq comparative analysis of transcripts quantified and also gene set enrichment analysis (GSEA). Data are from n=3 biologically independent replicates per group.

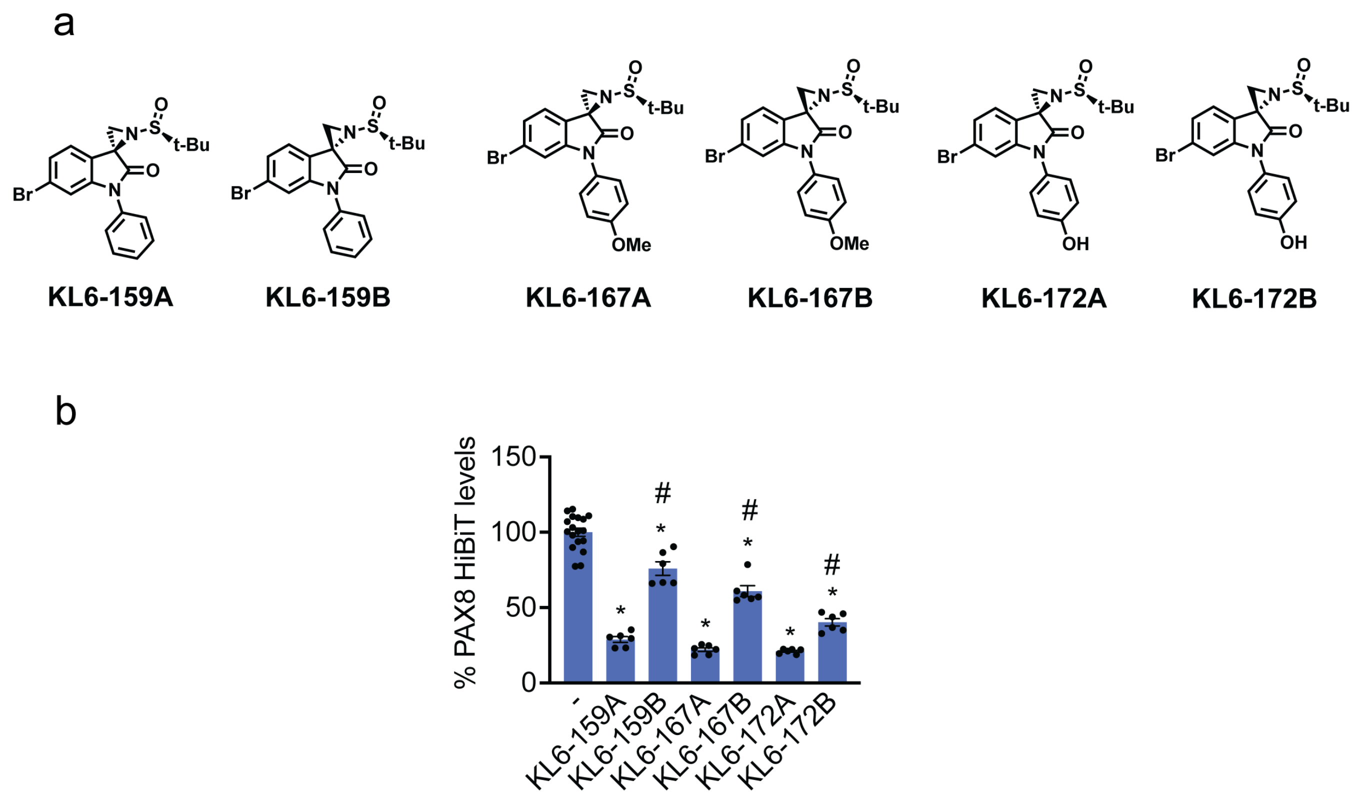


**Figure S1. Identification of KL6-159A as the lead stereoselective PAX8 destabilizer. (a)** Chemical structures of the stereochemical pairs corresponding to the initial screening hits KL6-159, KL6-167, and KL6-172. **(b)** Comparison of active compounds with their corresponding enantiomers in the HiBiT-PAX8 screening assay demonstrates stereoselective reduction of endogenous PAX8 by all three sulfinyl aziridine scaffolds, with KL6-159A exhibiting the greatest stereochemical discrimination and therefore selected for subsequent mechanistic studies. OVCAR3 HiBiT-PAX8-expressing cells were treated with DMSO vehicle or compounds (25 μM) for 24 h. Shown in **(b)** are individual replicate values and average ± sem. Statistical significance is shown as *$p<0.05$ compared to vehicle treated controls and #$p<0.05$ compared to the respective enantiomeric pair.

### Methods

#### Cell culture

HiBiT PAX8 CRISPR knock in with an OVCAR-3 background cells were acquired from Promega and cultured in RPMI with 25 mM HEPES (Gibco 22400-089) supplemented with 10% Fetal Bovine Serum (Corning 35-0101-CV). OVCAR-3 cells were obtained from UC Berkeley's Biosciences Divisional Services Cell Culture Facility and cultured in RPMI (Gibco 11875-093) supplemented with 10% Fetal Bovine Serum (Corning 35-0101-CV). OVISE cells were acquired from FujiFilm (JCRB1043) and cultured in RPMI (Gibco 11875-093) supplemented with 10% Fetal Bovine Serum (Corning 35-0101-CV). TOV-21G cells (CRL-3577) were acquired from American Type Culture Collection (ATCC) and cultured in a 1:1 mixture of MCDB 105 medium (Sigma-Aldrich M6395) containing a final concentration of 1.5 g/L sodium bicarbonate and Medium 199 (Thermo Fisher 11150067) containing a final concentration of 2.2 g/L sodium bicarbonate supplemented with 15% Fetal Bovine Serum (Corning 35-0101-CV). SKOV3 cells were obtained from UC Berkeley's Biosciences Divisional Services Cell Culture Facility and cultured in DMEM (Corning 10-0130CV) supplemented with 10% Fetal Bovine Serum (Corning 35-0101-CV). HEK293T cells were obtained from UC Berkeley's Biosciences Divisional Services Cell Culture Facility and cultured in DMEM (Corning 10-0130CV) supplemented with 10% Fetal Bovine Serum (Corning 35-0101-CV). All of the above cells were maintained at 37°C with 5% CO2.

#### Covalent Ligand Libraries

Structures of covalent ligands screened can be found in **Table S1.** Covalent ligand libraries were purchased from Enamine for compound names starting with "EN" or were synthesized as previously described for compound names starting with "KL" or "KLE" [1].

#### Screening and Testing of Covalent Ligands with HiBiT-PAX8 Cells

Covalent ligand screen and dose responses were conducted using the Nano-Glo HiBiT Lytic Detection System (Promega N3040) and CellTiter-Glo 2.0 Assay (Promega G9242). HiBiT PAX8 CRISPR knock in with an OVCAR-3 background cells were seeded at 25,000 cells per well in a 96-well plate (Corning 3917) in 99 µL of media for screening or 90 µL for dose-responses and were left to adhere overnight. For screening, cells were treated with 1 µL of DMSO 2.5 mM compound. For dose responses, cells were treated with 10 µL of compound diluted in media for the final concentration indicated. The Nano-Glo HiBiT Lytic Detection System and Cell Titer Glo 2.0 were used per the manufacturer's instructions (1:1 media to reagent). Plates were incubated in the dark for 10 minutes followed by using a Tecan Spark plate reader (30086376) to measure luminescence.

#### Western Blotting

Cells were seeded in 6-well plates (300,000 cells/well for OVCAR3, OVISE, and SKOV3 cells, 450,000 cells/well for TOV-21G) or 6 cm dishes (500,000 cells/dish for OVISE cells and 400,000 cells/well for HEK293T cells) and allowed to adhere overnight. For treating, the media was aspirated, replaced with media containing DMSO or compound, and treated for the indicated length of time. For harvesting, cells were then washed with PBS before lysing on the plate using RIPA buffer (Thermo Fisher Scientific 89900) containing 1 x protease inhibitor cocktail (Thermo Fisher Scientific A32955) and benzonase (Millipore 70746-3 at 25-29 unit/mL) diluted 1:1000 and incubating at 37°C for 5 minutes. Samples were collected and protein concentrations were normalized to 1-2 mg/mL using BCA Protein Assay (Pierce 23225). 4 x reducing Laemmli SDS sample loading buffer was added and samples were boiled for 5 minutes at 95°C and run on precast 4-20% Criterion TGX gels (Bio-Rad) in 1 x tris-glycine-sodium dodecyl sulfate (TGS) buffer (Bio-Rad 1610734). Proteins were resolved by sodium dodecyl sulfate-polyacrylamide gel electrophoresis (SDS-PAGE) and transferred to 0.2 µm nitrocellulose membrane using the Bio-Rad Trans-Blot Turbo Transfer system (Bio-Rad, 1704150 and 1704271). Membranes were blocked with 5% Bovine Serum Albumin (BSA) (bioWORLD 22070008-6) in Tris-buffered saline containing Tween 20 solution (TBST) (Thermo Fisher Scientific

J77500.K8) for 1 hour at room temperature. Blots were then incubated with primary antibody solution overnight at 4 °C with gentle rocking. Antibodies used in this study were PAX8 (CST 59019), vinculin (Invitrogen 14-9777-82), (Proteintech 60004-1-Ig), c-MYC (Abcam AB32072), FLAG (Sigma-Aldrich F1804), and LMB1 (Proteintech 66095-1-Ig) diluted 1:1000 in 5% BSA in TBST. Membranes were then washed 3 times with TBST, followed by 1 hour of incubation with a secondary antibody solution in the dark at room temperature. Secondary antibodies used in this study include IRDye680RD Goat anti-Mouse IgG (LicorBio, 926-68070) and IRDye800CW Goat anti-Rabbit IgG (LicorBio 926-32211) diluted 1:10,000 in 5% BSA in TBST solution. Before imaging, membranes were washed an additional 3 times with TBST. Western blots were visualized using an Odyssey DLx Imager (LICORbio) and ChemiDoc MP Imaging System (Bio-Rad).

**Tandem Mass Tag Based Global Quantitative Proteomics**

Cells were treated with DMSO vehicle or compound for 16 hours. For harvesting, cells were washed with PBS, harvested by scraping, and pelleted by centrifugation (1400 g, 4 min, 4 °C). Pellets were resuspended in PBS supplemented with protease inhibitor (Thermo Fisher Scientific A32955) and lysed by sonication. Protein concentrations were determined using the BCA assay (Pierce 23225). 25-100 µg of lysate was reduced, alkylated and digested with sequencing grade trypsin overnight. Individual samples were then labeled with isobaric tags using commercially available TMT10plex (Thermo Fisher Scientific 90110) kits, in accordance with the manufacturer's protocols. Tagged samples (20 µg per sample) were combined, dried using a vacuum concentrator at 30 °C, resuspended with 300 µL 0.1% trifluoroacetic acid (TFA) in water, and fractionated using high pH reversed-phase peptide fractionation kits (Thermo Fisher Scientific 84868) according to the manufacturer's protocol. Fractions were dried using a vacuum concentrator at 30 °C, resuspended with 50 µL 0.1% formic acid (FA) in water, and analyzed by LC-MS/MS as described below.
Mass spectrometry analysis was performed on an Orbitrap Eclipse Tribrid Mass Spectrometer with a High Field Asymmetric Waveform Ion Mobility (FAIMS Pro) Interface (Thermo Fisher Scientific) with an UltiMate 3000 Nano Flow Rapid Separation LCnano System (Thermo Fisher Scientific). Offline fractionated samples (5 µL aliquot of 50 µL sample) were injected via an autosampler (Thermo Fisher Scientific) onto a 5 µL sample loop, which was subsequently eluted onto an Acclaim PepMap 100 C18 HPLC column (75 µm × 50 cm, NanoViper). The peptides were separated at a flow rate of 0.3 µL/min using the following gradient: 2% buffer B (acetonitrile with 0.1% formic acid) in buffer A (95:5 water/acetonitrile, 0.1% formic acid) for 15 min, followed by a gradient from 2−20% buffer B from 15−115 min, 20−32% buffer B from 115−135 min, 32-95% buffer B from 135-136 min, held at 95% B from 136−160 min, 95% to 2% buffer B from 160−160.1 min, and then held at 2% buffer B until 180 min. The voltage applied to the nano-LC electrospray ionization source was 2.5 kV. Data were acquired through an MS1 master scan (Orbitrap analysis, resolution 120,000, 400−1600 *m/z*, RF lens 30%, ion transfer tube temperature 300 °C) with dynamic exclusion (repeat count 1, duration 60 sec). Data-dependent data acquisition comprised a full MS1 scan, followed by sequential MS2 scans based on 2.5 sec cycle times. FAIMS compensation voltages (CVs) of −40, −60, and −80 were applied. MS2 analysis consisted of a quadrupole isolation window of 0.7 *m/z* of the precursor ion followed by fragmentation via collision induced dissociation (CID) with 35% collision energy. Acquisition in the ion trap with scan rate set to turbo and then followed by a data dependent MS3 scan (10 SPS precursors, MS isolation window 0.7 *m/z*, MS2 isolation window 2 *m/z*, high energy collision dissociation (HCD), with a normalized collision energy of 55%, acquisition in the orbitrap with a resolution of 50,000).

Raw files were analyzed using the Chaparral Platform and SagePro. Trypsin cleavage specificity (cleavage at K, R, except if followed by P) allowed for up to 2 missed cleavages. Carbamidomethylation of cysteine residues (+57.02146) and TMT modification of peptide N-termini and lysine residues were set as static modification and methionine oxidation (+15.9949) was set as variable modification. MS1 tolerance was set to 10 ppm and MS2 tolerance to 300 ppm. Reporter-ion quantification was performed based on MS3 scans.

**Cellular Thermal Shift Assay**

OVCAR3 cells were treated with either DMSO control or 50 µM KL6-159A for 4 hours. Cells were harvested by scraping in PBS, centrifugation at 1,300 g for 5 minutes at 4 °C. Cells were resuspended in PBS containing 1 x protease inhibitor cocktail (Thermo Scientific A32955) and then aliquoted into eight 0.2 mL PCR strips with 85 µL per tube. PCR strips were designated a temperature via a gradient program on Bio-Rad's T100 Thermal cycler (Bio-Rad 1861096). Samples were heated at their respective temperatures (67 °C, 65 °C, 61.7 °C, 57 °C, 51.4 °C, 46.7 °C, 43.7 °C, 42 °C) for 3 minutes and then at 25 °C for 3 minutes. Immediately following, cells were snap-lysed using liquid nitrogen (3 freeze-thaw cycles). Cell debris, along with any precipitated and aggregated proteins, were removed by centrifugation at 20,000 g for 20 minutes at 4 °C. 81 µL of supernatant was transferred to new PCR strips, 27 µL of 4 x reducing Laemmli SDS sample loading buffer (Thermo Scientific J60015.AD) was added to each sample and boiled at 95 °C for 7 minutes. 14 µL of each sample was loaded per well into a precast 4-20% Criterion TGX gels (Bio Rad 5671095) in 1 x TGS buffer (Bio-Rad 1610772). Western blot prepared as stated above.

**Pure Protein Labeling with Covalent Alkyne Probe and Ligand Competition**

Recombinant full length PAX8 (CusaBio CSB-EP017494HU) was diluted to 0.5 µg/ 50 µL in PBS and aliquoted into PCR strips. Samples were either treated with either vehicle control (DMSO) or 150 µM KL6-159A and incubated at 37 °C for 30 minutes. Samples were then treated with either vehicle control or KL6-238A at the indicated concentrations and incubated at room temperature for 30 minutes. A master mix of the click reagents was prepared such that each replicate would receive 0.25 µL of Rhodamine Azide (5mM stock in DMSO, Click Chemistry Tools, AZ109-5), 1 µL of copper (II) sulfate (50mM stock in water, Sigma-Aldrich, 203165), 3 µL of TBTA (1.7 mM in 4:1 tBuOH/DMSO, TCI Chemicals, T2993) and 1 µL of Tris(2-carboxyethyl)phosphine, hydrochloride (TCEP) (50 mM in water freshly prepared, Strem Chemicals, 15-7400). Samples were gently vortexed and incubated at room temperature for 1 hour, after which 30 µL of 4 x reducing Laemmli SDS sample loading buffer was added to each replicate and boiled at 95 °C for 5 minutes. Samples were cooled to room temperature, 15 µL of sample per well was loaded into a precast 4-20% Criterion TGX gel (Bio Rad 5671095) and run in 1 x TGS buffer (Bio-Rad 1610734). Proteins were resolved by SDS/PAGE. Probe-labeled proteins were analyzed by in-gel fluorescence using the ChemiDoc MP Imaging system (Bio-Rad). Gels were silver stained using the Pierce Silver Stain Kit (Thermo Scientific 24612) per manufacturer instructions and imaged on the ChemiDoc MP Imaging System (Bio-Rad).

**Pulldown of PAX8 with Covalent Alkyne Probe and Ligand Competition for Western Blotting Detection**

Six 10 cm dishes with 200,000 OVISE cells each were seeded and allowed to adhere overnight. Cells were treated with either DMSO or 100 µM KL6-159A for 1 hour followed by treatment with either DMSO or 25 µM KL6-283A for 2 hours. Cells were washed with PBS, harvested by scraping, and pelleted by centrifugation. Pellets were resuspended in PBS with protease inhibitor (Thermo Scientific A32955) and lysed on ice by sonication (4 x 15s at 15% amplitude). Lysate was clarified by centrifugation at 4 °C for 10 minutes.

Lysate protein concentrations were measured using BCA Protein Assay (Pierce 23225) and normalized to 135 µg/mL in 500 µL of PBS with protease inhibitor cocktail (Thermo Scientific A32955). A master mix of the click reagents was prepared such that each replicate would receive 4 µL of biotin picolyl azide (10 mM stock in DMSO, Sigma-Aldrich 900912), 4 µL of copper (II) sulfate (50 mM stock in water, Sigma-Aldrich, 203165), 12 µL of TBTA (1.7 mM in 4:1 tBuOH/DMSO, TCI Chemicals T2993) and 10 µL of TCEP (50mM in water freshly prepared, Strem Chemicals 15-7400). Samples were vortexed and incubated on a rotator at room temperature for 1 hour. Proteins were precipitated with acetonitrile and pelleted by centrifugation (6500 g) at room temperature for 5 minutes. The supernatant was removed, 500 µL of cold methanol was added, and samples underwent three cold methanol washes with centrifugation to pellet proteins and sonication for resuspension.

Pellets were redissolved in 200 µL PBS with 1.2% SDS (w/v) and heated at 90 °C for 5 minutes. 5 µL of the sample was removed and saved for input (diluted to 60 µL with PBS and added 20 µL of 4 x reducing Laemmli SDS sample loading buffer). 55 µL (per sample) of streptavidin-agarose beads (ThermoFisher

20353) were washed with PBS using a Micro-Bio Spin Column (Bio-Rad 7326204) and vacuum manifold. 500 µL of PBS was added to the dissolved proteins, and then the washed beads were added to each sample using two washes of 250 µL of PBS. Samples were incubated on a rotator at 4 °C overnight. Samples were then put in a 37 °C bath to redissolve SDS for 5 minutes, and then beads were pelleted by centrifugation at 1400 g for 5 min. The supernatant was removed, and beads were washed with 0.2% SDS in PBS (w/v) for 10 minutes on a rotator at room temperature. The supernatant was removed, and pelleted beads were moved to Micro-Bio Spin Columns using two washes of PBS. Beads were washed on a vacuum manifold three times with PBS and then three times with water. The beads were then moved to screw cap Eppendorf tubes with PBS, centrifuged, and the supernatant removed. 30 µL of 1 x reducing Laemmli SDS sample loading buffer was added to each sample and then boiled at 95 °C for 15 minutes to elute proteins from beads. Samples and corresponding inputs were run on precast 4- 20% Criterion TGX gels (Bio Rad 5671095) in 1 x TGS buffer (Bio-Rad 1610734). Western blots were visualized using an Odyssey DLx Imager (LICORbio). Western blot prepared as stated above.

**Protein Half-Life Analysis with Treatment of Cycloheximide**
OVISE cells were seeded at 250,000 cells per well in 6-well plates and allowed to adhere overnight. Cells were treated with either 100 µg/mL cycloheximide or cotreated with 100 µg/mL cycloheximide and 50 µM KL6-159A for the indicated time points. Cells were washed with PBS and lysed off the plate with 60 µL RIPA buffer with 1 x protease inhibitor (Thermo Scientific A32955) and 1 x benzonase (Millipore 70746-3). 60 µL of lysate per sample was collected into PCR strips. Lysates were normalized to 0.9 µg/ µL using BCA Protein Assay (Pierce 23225) in 60 µL. 20 µL of 4 x reducing Laemmli SDS sample loading buffer (Thermo Scientific J60015.AD) was added to each sample and boiled at 95 °C for 5 minutes. 15 µL of each sample was loaded per well into a precast 4-20% Criterion TGX gels (Bio Rad 5671095) in 1 x TGS buffer (Bio-Rad 1610772). Western blot prepared as stated above.

**Generating Stable Cell Lines via Lentivirus for Firefly Luciferase Under a PAX8 Promoter**
A plasmid for firefly luciferase under a PAX8 promoter was customized and purchased from vector builder. psPAX2 (Addgene 12260) and pMD2.G (Addgene 12259) were transfected into HEK293T cells using Lipofectamine 2000 (ThermoFisher 11668027) in Opti-MEM (Gibco 31985062). The virus-containing medium was collected and filtered (0.45 µm PES) after 48 hours. The virus was then used to infect OVISE cells with a 1:1000 dilution of Polybrene (Sigma-Aldrich TR-1003-G). After 48 hours, infected cells were selected with 0.2 µg/mL puromycin (Abcam ab141453). After 4 days of the selection, cells were recovered by replacing puromycin-containing media with fresh media.

**Luciferase Reporter Assay**
OVISE cells stably expressing luciferase under a PAX8 promoter were seeded at 20,000 cells per well in 100 µL in 96-well plates. Cells were allowed to adhere overnight at 37 °C with 5% $CO_2$ before treating. Bright-Glo luciferase assay (Promega E2610) was used according to the manufacturer's protocol for luminescent readout. Firefly luminescence was read on a Tecan Spark plate reader. Background luminescence levels were subtracted using a blank control, and then firefly luminescence signal was calculated for each well.

**Activity-Based Protein Profiling Cysteine Chemoproteomics by Isotopically Labeled Desthiobiotin Azide**
Fifteen 15 cm plates of OVISE cells were treated with either vehicle control (DMSO) or 50 µM KL6-159A for 2 hours. Cells were harvested by washing with PBS, scraping in PBS, and centrifugation at 300 g for 4 minutes. Cells were resuspended in PBS with 1 x protease inhibitor (Thermo Scientific A32955) and then lysed by sonication at 15% amplitude for 15 seconds 6 times. Lysates were clarified by centrifugation at 10,000 g for 10 minutes at 4 °C. Lysates were normalized to 2 mg/mL using BCA Protein Assay (Pierce 23225) and 3 x 1 mL was aliquoted into microcentrifuge tubes per condition. N-hexyl-5-ynyl-2iodo-acetamide

(Iodoacetamide alkyne, IA alkyne) (Oakwood Chemicals 095226) in DMSO was added for a concentration of 200 µM and incubated at room temperature for 1 hour with end-over-end mixing. Two master mixes with 165 µL of $CuSO_4$ (50mM stock in water, Sigma-Aldrich 203165), 165 µL of TCEP (50 mM in water freshly prepared, Strem Chemicals 15-7400), and TBTA (1.7 mM in 4:1 tBuOH/DMSO, TCI Chemicals T2993) were made. 4 mg of isotopically labeled desthoibiotin azide probes (light and heavy) were dissolved in 160 µL of DMSO. The light probe was transferred into one of the master mixes and the heavy probe into the other master mix. 120 µL of the light master mix was transferred into each replicate of the DMSO treated samples. 120 µL of the heavy master mix was transferred into each replicate of the 50 µM KL6-159A treated samples. The samples were incubated at room temperature for 2 hours with end over end mixing. After isotopic labeling, samples were combined such that one DMSO sample and one compound treated sample were combined and precipitated with acetonitrile at -20 °C overnight.

Protein precipitates were pelleted by centrifugation at 3,500 rpm for 10 minutes. The supernatant was removed and the proteins were washed three times by resuspending in cold methanol by sonication (3 x 15% amplitude for 15 seconds) and pelleting by centrifugation (3,500 rpm for 10 minutes). Pelleted were then dissolved in 600 µL 8M urea in 0.1 M triethylammonium bicarbonate (TEAB) by sonication (3 x 15% amplitude for 15 seconds). 1800 µL of 0.1 M TEAB, 2400 µL of 0.2% NP-40 (Thermo Scientific 85124) diluted in PBS, and 200 µL of high-capacity streptavidin agarose resin (Thermo Scientific 20361) were added to each sample. Samples were incubated at room temperature for 1 hour with end over end mixing. Streptavidin beads were pelleted by centrifugation at 1,000 g for 1 minute and supernatant was removed. Streptavidin beads were washed with 3 times with 1 mL of 0.1% NP-40 (Thermo Scientific 85124) diluted in PBS, 3 times with 1 mL of PBS and 3 times with 1 mL of MilliQ water. Beads were resuspended in 600 µL of 8M urea in 0.1M TEAB. 8 µL of 0.5 mg/mL sequencing grade modified trypsin (Promega V5111) resuspended in trypsin resuspension buffer (Promega V542A) was added to each sample. Samples were incubated overnight at 37 °C and 250 rpm.

Samples were diluted with 800 µL 0.1% NP-40 (Thermo Scientific 85124) diluted in PBS and transferred to a microspin column (Bio-Rad 7326204) equipped with a vacuum manifold. Beads were washed with 1 mL 3 times with 0.1 %NP-40 (Thermo Scientific 85124) diluted in PBS, PBS, and MilliQ water. The microspin columns were transferred to 2 mL microcentrifuge tubes. 500 µL of 0.1% formic acid in 50% acetonitrile were added to the microspin columns and then the microspin columns were capped and the beads were incubated for 5 minutes. The samples were centrifuged at 1,400 g for 3 minutes. The elution step was repeated two more times, and the peptides were dried with a vacuum concentrator. The peptides were resuspended in 300 µL 0.1% TFA and fractionated according to Pierce high pH reverse phase peptide fractionation kit (Thermo Scientific 84868). Fractions were dried with a vacuum concentrator, resuspended in 25 µL of 0.1% formic acid in water and centrifuged at 20,000 g for 5 minutes. Supernatant was transferred to LC-MS vials with low volume inserts.

Mass spectrometry analysis was performed on an Orbitrap Eclipse Tribrid Mass Spectrometer with a High Field Asymmetric Waveform Ion Mobility (FAIMS Pro) Interface (Thermo Fisher Scientific) with an UltiMate 3000 Nano Flow Rapid Separation LCnano System (Thermo Fisher Scientific). Offline fractionated samples (5 µL aliquot of 25 µL sample) were injected via an autosampler (Thermo Fisher Scientific) onto a 5 µL sample loop, which was subsequently eluted onto an Acclaim PepMap 100 C18 HPLC column (75 µm × 50 cm, NanoViper). The peptides were separated at a flow rate of 0.3 µL/min using the following gradient: 2% buffer B (acetonitrile with 0.1% formic acid) in buffer A (95:5 water/acetonitrile, 0.1% formic acid) for 5 min, followed by a gradient from 2−40% buffer B from 5−159 min, 40−95% buffer B from 159−160 min, held at 95% B from 160−179 min, 95% to 2% buffer B from 179−180 min, and then 2% buffer B from 180−200 min. The voltage applied to the nano-LC electrospray ionization source was 2.1 kV. Data were acquired through an MS1 master scan (Orbitrap analysis, resolution 120,000, 400−1800 *m/z*, RF lens 30%, heated capillary temperature 250 °C) with dynamic exclusion (repeat count 1, duration 60 sec). Data-dependent data acquisition comprised a full MS1 scan, followed by sequential MS2 scans based on 2 sec cycle times. FAIMS compensation voltages (CVs) of −35, −45, and −55 were applied. MS2 analysis consisted of a quadrupole isolation window of 0.7 *m/z* of the precursor ion followed by a higher energy collision dissociation (HCD) energy of 38% with an orbitrap resolution of 50,000.

The obtained raw-data was analyzed using FragPipe (v24) using an adjusted ABPP-isoDTB[2] workflow using the whole human proteome (uniport, downloaded 01/30/2025) as search space and applying the following MSFragger[3,4]settings: Precursor mass tolerance: +/-20 ppm; Fragment mass tolerance: +/-20 ppm; Mass calibration & parameter optimization enabled; Isotope error: 0/1/2; Enzyme: Trypsin (cuts after K & R, no cut before P), Peptide length: 6-50 AA; Peptide mass range: 500-5000 Da; Variable modifications: Oxidation (M, 15.9949 Da, up to 2 x), Acetylation (any protein N-term, 42.0106 Da); Carbamidomethylation of C was set as fixed modification (57.02146 Da). Additionally, cysteine modification with heavy/light isoDTB tags was set as variable modification (+561.3387/567.3462 Da). Rescoring via MSBooster[5] was enabled and validation was performed via Percolator & ProteinProphet with a 1% FDR. IonQuant[6,7] was used for quantification, using light & heavy isoDTB-tags as labels with Re-quantify and MBR (FDR of 1%) enabled.

**Generating PAX8-FLAG Wild-Type and Mutants**

Human PAX8-FLAG constructs were customized purchased from vector builder. The customized construct was in a pLV backbone and PAX8 had a C-terminal DYKDDDDK (FLAG) tag.

HEK293T cells were seeded at 400,000 cells per well in 6-well plates and cells were allowed to adhere overnight. Cells were transfected with 1.25 µg of plasmid per well using lipofectamine 3000 (Invitrogen L3000008) for 24 hours according to manufacturer protocol. Media containing transfection reagents was replaced with fresh media containing either DMSO control or 50 µM KL6-159A for 4 hours. Cells were harvested by aspirating media, washing with PBS and then scraping with PBS. Cells were pelleted by centrifugation at 300 g for 4 minutes at 4 °C. The supernatant was removed and cells were resuspended in 50 µL hypotonic lysis buffer (10 mM EPPS pH8, 1.5 mL $MgCl_2$, and 10 mM KCl) with 1 x protease inhibitor (Med Chem express HY-K0010) and incubated on ice for 15 minutes. 2.5 µL of 10% NP-40 was added and samples were vortexed gently and centrifuged at 3000 rpm for 10 minutes at 4 °C. The supernatant was removed as the cytoplasmic fraction. The remaining pellet was lysed with RIPA buffer (Thermo Scientific 89900) containing protease inhibitor cocktail (Thermo Scientific A32955) and Benzonase (Millipore 70746-3 at 25-29 unit/mL) at 37°C for 5 minutes. To the lysate, 17.5 µL of 4 x reducing Laemmli SDS sample loading buffer was added and samples were boiled for 5 minutes at 95 °C and run on precast 4-20% Criterion TGX gels (Bio-Rad) in 1 x TGS buffer (Bio-Rad 1610734). Proteins were resolved by SDS/PAGE and transferred to 0.2 µm nitrocellulose membrane using the Bio-Rad Trans-Blot Turbo Transfer system (Bio-Rad, 1704150 and 1704271). Membranes were blocked with 5% Bovine Serum Albumin (BSA) (bioWORLD 22070008-6) in Tris- buffered saline containing Tween 20 solution (TBST) (ThermoFisher J77500.K8) for 1 hour at room temperature. Blots were then incubated with primary antibody solution overnight at 4°C with gentle rocking. Antibodies used in this study were FLAG (Sigma-Aldrich F1804), LMB1 (Proteintech 66095-1-Ig) and vinculin (Invitrogen 14-9777-82) were diluted 1:1000 in 5% BSA in TBST. Membranes were then washed 3 times with TBST, followed by 1 hour of incubation with a secondary antibody solution in the dark at room temperature. Secondary antibodies used in this study include IRDye680RD Goat anti-Mouse IgG (LicorBio, 926-68070) and IRDye800CW Goat anti-Rabbit IgG (LicorBio 926-32211) diluted 1:10,000 in 5% BSA in TBST solution. Before imaging, membranes were washed an additional 3 times with TBST. Western blots were visualized using an Odyssey DLx Imager (LICORbio) and ChemiDoc MP Imaging System (Bio-Rad).

**RNA Extraction and Sequencing**

OVISE cells were seeded at 200,000 cells per well in 6-well plates and allowed to adhere overnight. Cells were treated with either DMSO control, 50 µM KL6-159A or 50 µM KL6-159B for 16 hours. Total RNA was extracted by aspirating the media, washing the cells with PBS, and then using RNeasy plus micro kit (Qiagen 74034) according to manufacturer protocol.

RNA quality control and sequencing was performed by Signios Biosciences. FASTQ raw files were aligned and quantified using Kallisto[8] and filtered for >10 transcripts per million (tpm) for all replicates of at least one condition followed by differential gene expression analysis using DESeq2[9] and a threshold of $|log_2(FoldChange)| > 1$ and an adjusted p-value < 0.05 was applied to identify differentially expressed genes. fGSEA analysis[10] was performed to identify significantly regulated MSigDB Hallmark gene sets[11]. Significantly downregulated genes were additionally compared to the ChEA 2022 TF-target dataset using Enrichr[12,13].

### Synthetic Methods and Characterization

#### General Procedures

All reactions were performed in flame- or oven-dried glassware under a positive pressure of nitrogen or argon, unless otherwise noted. Air- and moisture-sensitive liquids were transferred via syringe. Volatile solvents were removed under reduced pressure using rotary evaporation below 35 °C.

Analytical and preparative thin-layer chromatography (TLC) were performed using glass plates pre-coated with silica gel (0.25-mm, 60-Å pore size, Merck TLC Silicagel 60 F254) impregnated with a fluorescent indicator (254 nm). TLC plates were visualized by exposure to ultraviolet light (UV) and then were stained by submersion in an ethanolic p-anisaldehyde solution, ceric ammonium molybdate solution, or potassium permanganate solution followed by brief heating on a hot plate. Flash column chromatography was performed with HC Duo Silica columns purchased from Biotage (Biotage FSUD-0443). All HRMS data was collected by ESI-MS on a Thermo Q Exactive Plus with a mobile phase of 100% methanol. Synthetic procedures used to prepare oxindole sulfonyl and sulfinyl aziridines were adapted from the prior methods of Hajra and co-workers[14].

**General procedure A: Synthesis of *N*-aryl 6-Bromoisatin**. To a suspension of 6-Bromoisatin (1 equiv.), $Cu(OAc)_2 \cdot H_2O$ (1 equiv.), and $ArB(OH)_2$ (1.2 equiv.) in MeCN (0.16 M) under air $NEt_3$ (2 equiv.) was added upon which the reaction mixture turned dark purple. The reaction mixture was stirred at ambient temperature (ca. 20 °C) for 16 h or until starting material was consumed. The mixture was then filtered through a silica plug to remove Cu salts, the plug was washed with EtOAc, then the filtrate was concentrated *in vacuo*. The dark brown residue was purified by flash column chromatography on silica gel (5% → 30% EtOAc–hexanes) where fractions were collected once the orange-colored product began eluting to yield arylated 6-bromoisatins as orange solids.

**General procedure B: Synthesis of *N*-sulfinyl spiroaziridines** (KL6-159A, KL6-159B, KL6-167A, KL6-167B, KL6-172A, KL6-172B, KL6-294A, KL6-294B). i. To a mixture of *N*-substituted 6-Bromoisatin (90.5 mg, 0.300 mmol, 1 equiv.) and (S)-tert-butanesulfinamide (45.4 mg, 0.374 mmol, 1.25 equiv.) in anhydrous DCM (1.2 mL) at 20 °C $Ti(OiPr)_4$ (0.599 mmol, 2 equiv.) was added upon which the color of the mixture immediately darkened. The reaction vessel was sealed with Teflon tape, and the reaction mixture was heated to 50 °C for 6 hours with stirring. The reaction was cooled to room temperature and diluted with EtOAc (3 mL). Excess $Ti(O\text{-}iPr)_4$ was quenched by the addition of saturated *aq.* $NaHCO_3$ (2 mL), and the emulsion formed was vigorously stirred for 10 minutes at which point a white aq. suspension of $TiO_2$ separated. The mixture was filtered through a pad of celite and rinsed with EtOAc until colorless. The orange filtrate was washed with brine (10 mL), dried over $Na_2SO_4$, and concentrated *in vacuo* to yield the crude sulfinimine as a red oil that solidified upon standing. The crude sulfinimine was used immediately without further purification.

ii. A mixture of $Me_3SOI$ (71.8 mg, 0.326 mmol, 2 equiv.), $Cs_2CO_3$ (106 mg, 0.326 mmol, 2 equiv.), and powdered, activated 4Å molecular sieves (ca. 75 mg) in anhydrous MeCN (1 mL) was heated to 50 °C and stirred for 0.5 hours. After cooling to 20 °C, a solution of the crude sulfinimine (0.250 mmol, 1 equiv.) in anhydrous MeCN (1 mL) was added, and additional MeCN (2 x 0.5 mL) was used for quantitative transfer. The orange-colored reaction mixture was stirred at 20 °C for 4 hours during which the orange color gradually turned beige. The suspension was filtered through a pad of celite, rinsing with EtOAc, to remove the 4Å molecular sieves. The filtrate was then washed with brine (5 mL), dried over Na2SO4, and concentrated in vacuo. The crude residue was purified b flash column chromatography on silica gel (0% → 40% EtOAc–hexanes) to yield (S,S)- and (S,R)-spiroaziridines respectively as white or yellow solids. The combined yield of the two diastereomers ranged from 40% to 60%, and the d.r. ranged from 3:1 to 4:1 (S,S):(S,R).

KL6-294A and KL6-294B were synthesized using General Procedure B except using (R)-tert butanesulfinamide instead of (S)-tert-butanesulfinamide.

**KL6-159A:**
**$^1$H NMR** (500 MHz, $CDCl_3$) δ 7.60 (d, *J* = 8.2 Hz, 1H), 7.58 – 7.53 (m, 2H), 7.47 – 7.43 (m, 1H), 7.43 – 7.40 (m, 2H), 7.24 (dd, *J* = 8.2, 1.8 Hz, 1H), 7.03 (d, *J* = 1.7 Hz, 1H), 3.46 (s, 1H), 2.87 (s, 1H), 1.33 (s, 9H).
**$^{13}C\{^1H\}$ NMR** (126 MHz, $CDCl_3$) δ 171.0, 146.6, 133.7, 130.0, 128.8, 126.9, 126.7, 126.3, 123.5, 120.3, 113.5, 58.9, 44.2, 31.7, 22.6.
**HRMS** (ESI) *m/z*: $[M+H]^+$ Calcd for C19H20BrN2O2S 419.0423; Found 419.0419

**KL6-159B:**
**$^1$H NMR** (500 MHz, $CDCl_3$) δ 7.57 – 7.51 (m, 2H), 7.47 – 7.40 (m, 3H), 7.27 (s, 1H), 7.25 (d, *J* = 1.7 Hz, 1H), 7.06 (d, *J* = 1.7 Hz, 1H), 7.03 (d, *J* = 8.0 Hz, 1H), 3.44 (s, 1H), 2.79 (s, 1H), 1.27 (s, 9H).
**HRMS** (ESI) *m/z*: $[M+H]^+$ Calcd for C19H20BrN2O2S 419.0423; Found 419.0421

**KL6-294A:**
**$^1$H NMR** and **$^{13}C\{^1H\}$ NMR** data matched that of KL6-159A.
**HRMS** (ESI) *m/z*: $[M+H]^+$ Calcd for C19H20BrN2O2S 419.0423; Found 419.0422

**KL6-294B:**
**$^1$H NMR** and **$^{13}C\{^1H\}$ NMR** data matched that of KL6-159B.
**HRMS** (ESI) *m/z*: $[M+H]^+$ Calcd for C19H20BrN2O2S 419.0423; Found 419.0418

**KL6-283A:**
**$^1$H NMR** (500 MHz, $CDCl_3$) δ 7.52 (d, *J* = 8.1 Hz, 1H), 7.22 – 7.13 (m, 2H), 3.85 (td, *J* = 7.2, 4.0 Hz, 2H), 3.37 (s, 1H), 2.77 (s, 1H), 2.30 (td, *J* = 6.9, 2.7 Hz, 2H), 1.93 (p, *J* = 7.0 Hz, 2H), 1.30 (s, 9H).
**$^{13}C\{^1H\}$ NMR** (151 MHz, $CDCl_3$) δ 171.7, 146.0, 126.8, 125.6, 123.4, 120.5, 112.5, 82.9, 69.7, 58.7, 43.9, 39.7, 31.2, 26.2, 22.5, 16.3.
**HRMS** (ESI) *m/z*: $[M+H]^+$ Calcd for C18H22BrN2O2S 409.0580; Found 409.0573

**KL6-167A:**
**$^1$H NMR** (500 MHz, $CDCl_3$) δ 7.58 (d, *J* = 8.2 Hz, 1H), 7.32 – 7.30 (m, 2H), 7.21 (dd, *J* = 8.2, 1.8 Hz, 1H), 7.10 – 7.01 (m, 2H), 7.00 – 6.93 (m, 1H), 3.87 (s, 3H), 3.44 (s, 1H), 2.86 (s, 1H), 1.33 (s, 9H).
**HRMS** (ESI) *m/z*: $[M+H]^+$ Calcd for C20H22BrN2O3S 449.0529; Found 449.0524

**KL6-167B:**
**$^1$H NMR** (500 MHz, $CDCl_3$) δ 7.35 – 7.28 (m, 2H), 7.24 (dd, *J* = 7.9, 1.7 Hz, 1H), 7.10 – 6.95 (m, 4H), 3.86 (s, 3H), 3.42 (s, 1H), 2.78 (s, 1H), 1.26 (s, 9H).
**HRMS** (ESI) *m/z*: $[M+H]^+$ Calcd for C20H22BrN2O3S 449.0529; Found 449.0525

**KL6-172A:**
**$^1$H NMR** (700 MHz, $CDCl_3$) δ 7.60 (d, *J* = 8.2 Hz, 1H), 7.25 (dd, *J* = 8.2, 1.8 Hz, 1H), 7.24 – 7.20 (m, 2H), 6.96 (d, *J* = 1.8 Hz, 1H), 6.94 – 6.90 (m, 2H), 6.15 (s, 1H), 3.49 (s, 1H), 2.92 (s, 1H), 1.36 (s, 9H).
**HRMS** (ESI) *m/z*: $[M+H]^+$ Calcd for C19H20BrN2O3S 435.0373; Found 435.0369

**KL6-172B:**
**$^1$H NMR** (700 MHz, $CDCl_3$) δ 7.24 (dd, *J* = 8.6, 2.2 Hz, 1H), 7.23 – 7.20 (m, 2H), 7.01 (d, *J* = 7.9 Hz, 1H),

6.99 (d, $J$ = 1.6 Hz, 1H), 6.97 – 6.95 (m, 2H), 3.43 (s, 1H), 2.79 (s, 1H), 1.28 (s, 9H).
**HRMS** (ESI) *m/z*: $[M+H]^+$ Calcd for C19H20BrN2O3S 435.0373; Found 435.0368

**KL6-159A**

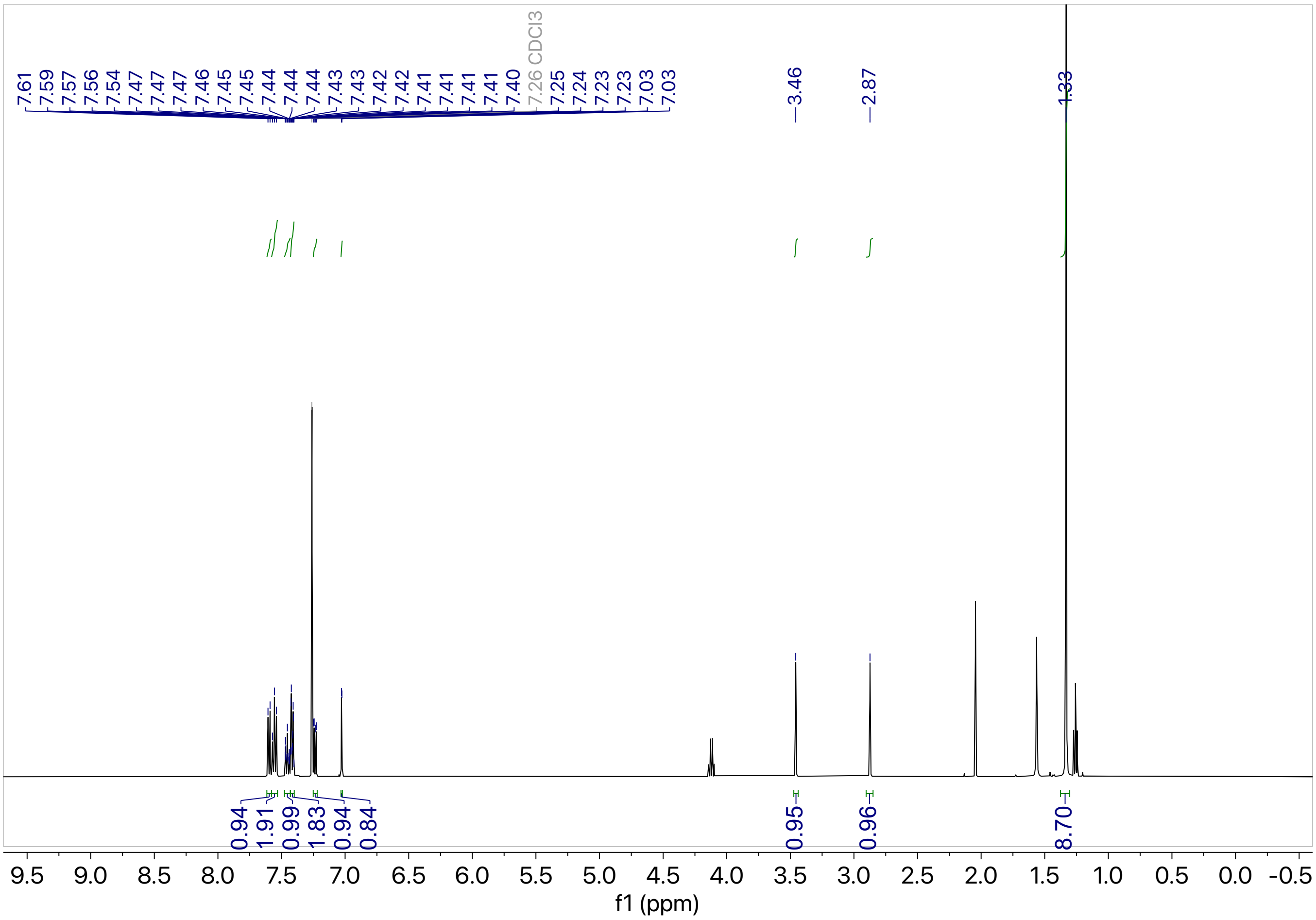

7.61
7.59
7.57
7.56
7.54
7.47
7.47
7.47
7.46
7.45
7.45
7.44
7.44
7.44
7.43
7.43
7.42
7.42
7.41
7.41
7.41
7.41
7.40
7.26 CDCl3
7.25
7.24
7.23
7.23
7.03
7.03
3.46
2.87
1.33
0.94
1.91
0.99
1.83
0.94
0.84
0.95
0.96
8.70
f1 (ppm)


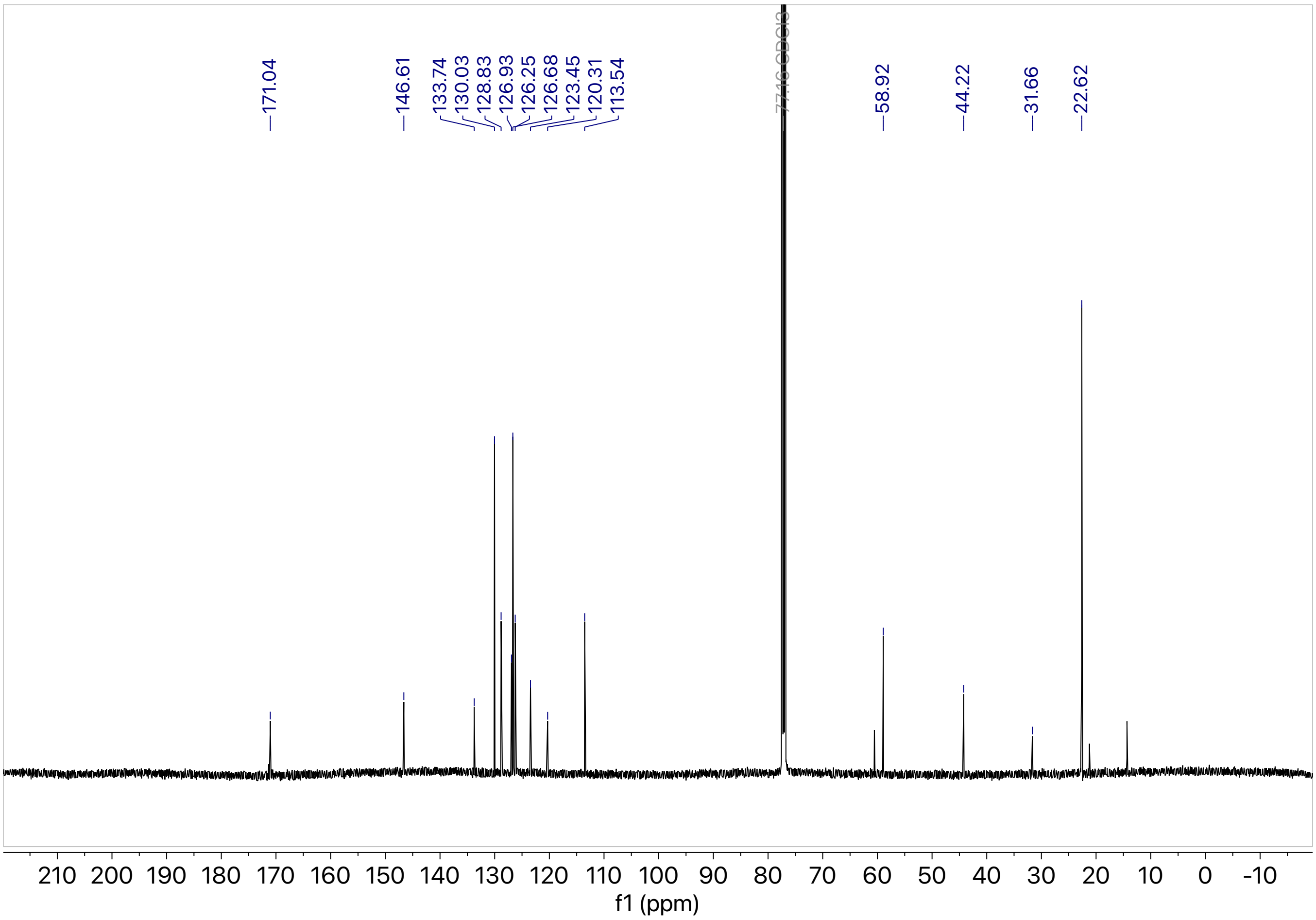

171.04
146.61
133.74
130.03
128.83
126.93
126.25
126.68
123.45
120.31
113.54
77.16 CDCl3
58.92
44.22
31.66
22.62
f1 (ppm)

**KL6-159B**

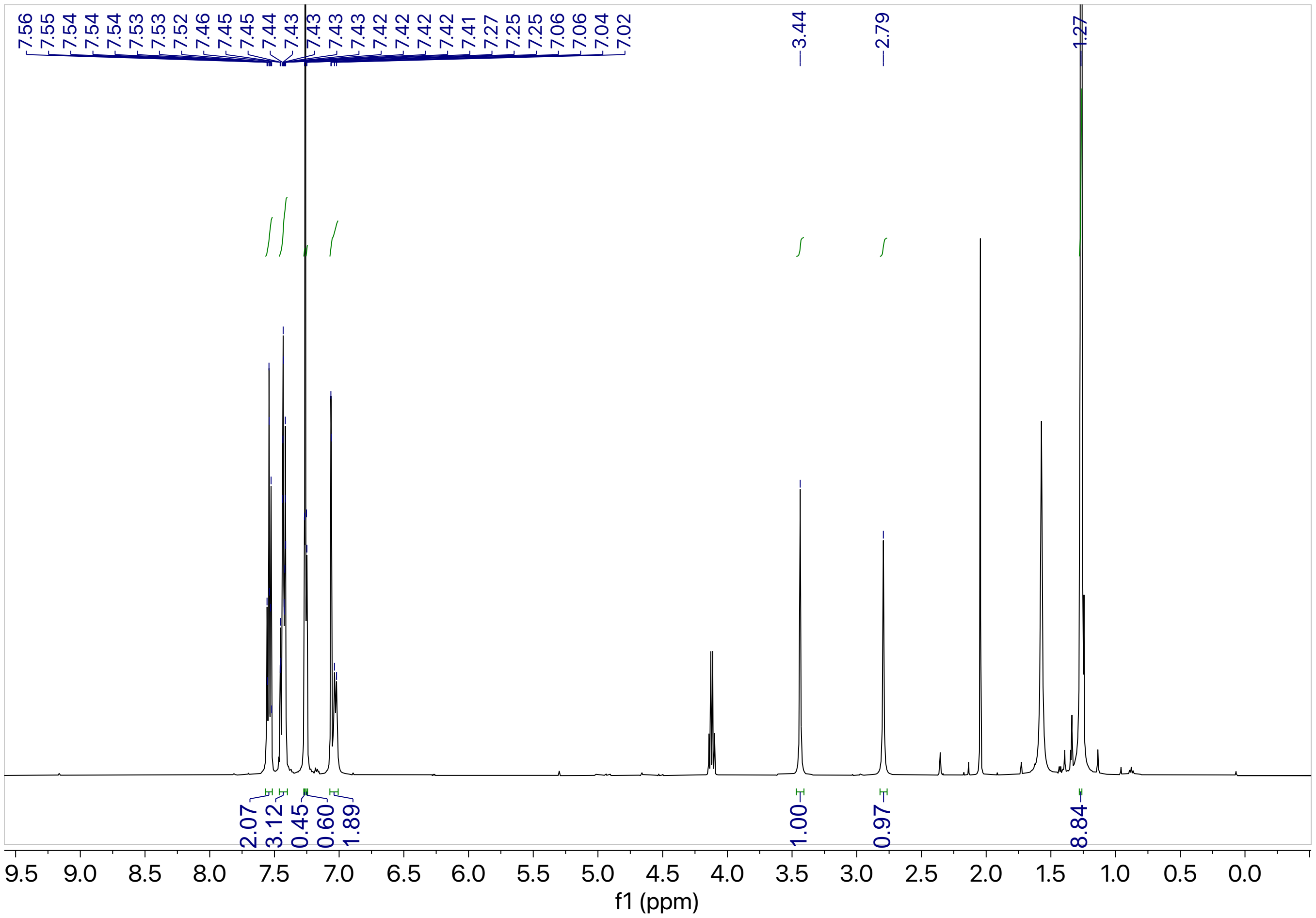

7.56
7.55
7.54
7.54
7.54
7.53
7.53
7.52
7.46
7.45
7.45
7.44
7.43
7.43
7.43
7.43
7.42
7.42
7.42
7.42
7.41
7.27
7.25
7.25
7.06
7.06
7.04
7.02
3.44
2.79
1.27
2.07
3.12
0.45
0.60
1.89
1.00
0.97
8.84
9.5 9.0 8.5 8.0 7.5 7.0 6.5 6.0 5.5 5.0 4.5 4.0 3.5 3.0 2.5 2.0 1.5 1.0 0.5 0.0
f1 (ppm)

**KL6-283A**

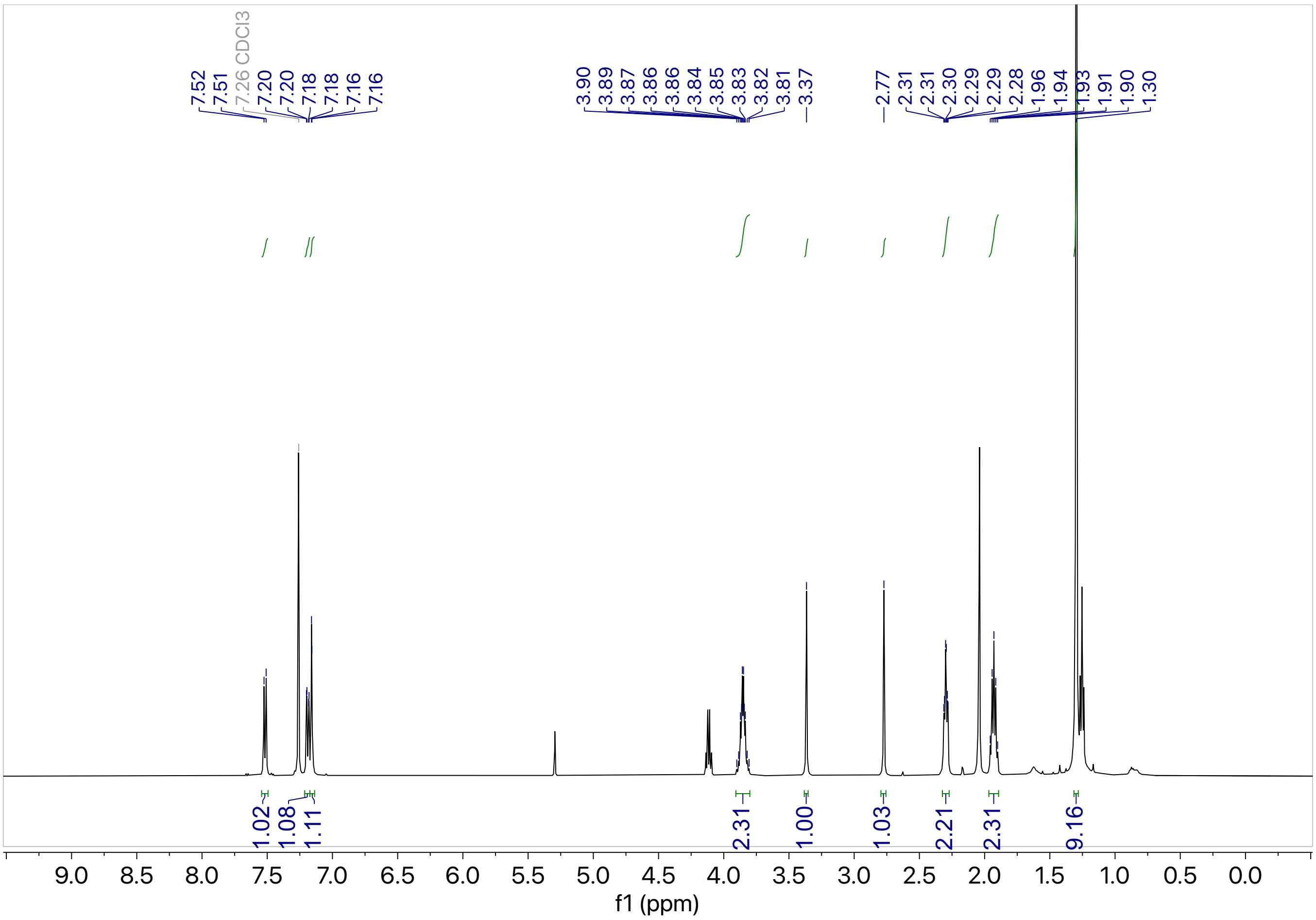

7.52
7.51
7.26 CDCl3
7.20
7.20
7.18
7.18
7.16
7.16
3.90
3.89
3.87
3.86
3.86
3.84
3.85
3.83
3.82
3.81
3.37
2.77
2.31
2.31
2.30
2.29
2.29
2.28
1.96
1.94
1.93
1.91
1.90
1.30
1.02
1.08
1.11
2.31
1.00
1.03
2.21
2.31
9.16
9.0 8.5 8.0 7.5 7.0 6.5 6.0 5.5 5.0 4.5 4.0 3.5 3.0 2.5 2.0 1.5 1.0 0.5 0.0
f1 (ppm)


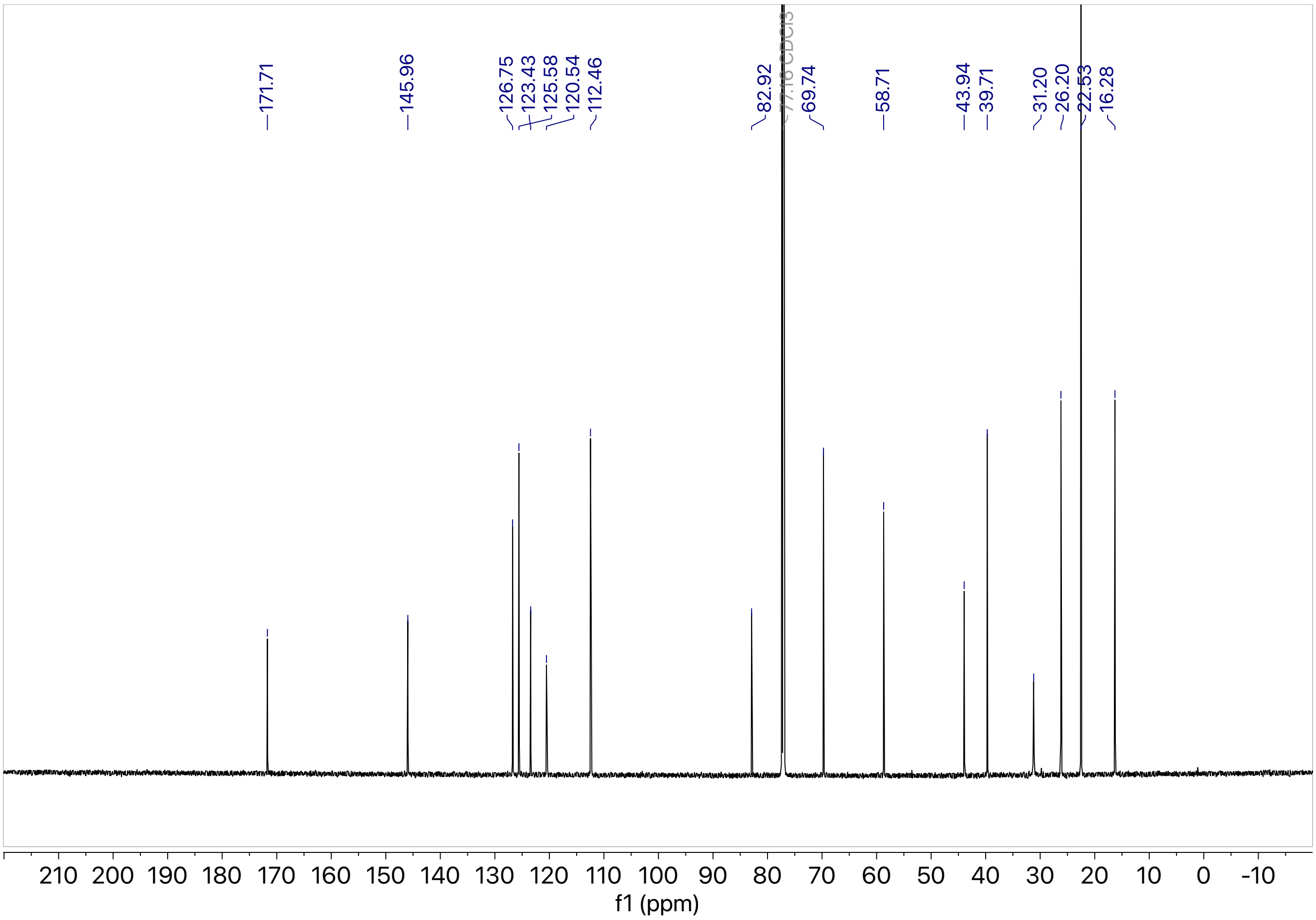

171.71
145.96
126.75
123.43
125.58
120.54
112.46
82.92
77.16 CDCl3
69.74
58.71
43.94
39.71
31.20
26.20
22.53
16.28
210 200 190 180 170 160 150 140 130 120 110 100 90 80 70 60 50 40 30 20 10 0 -10
f1 (ppm)

**KL6-167A**

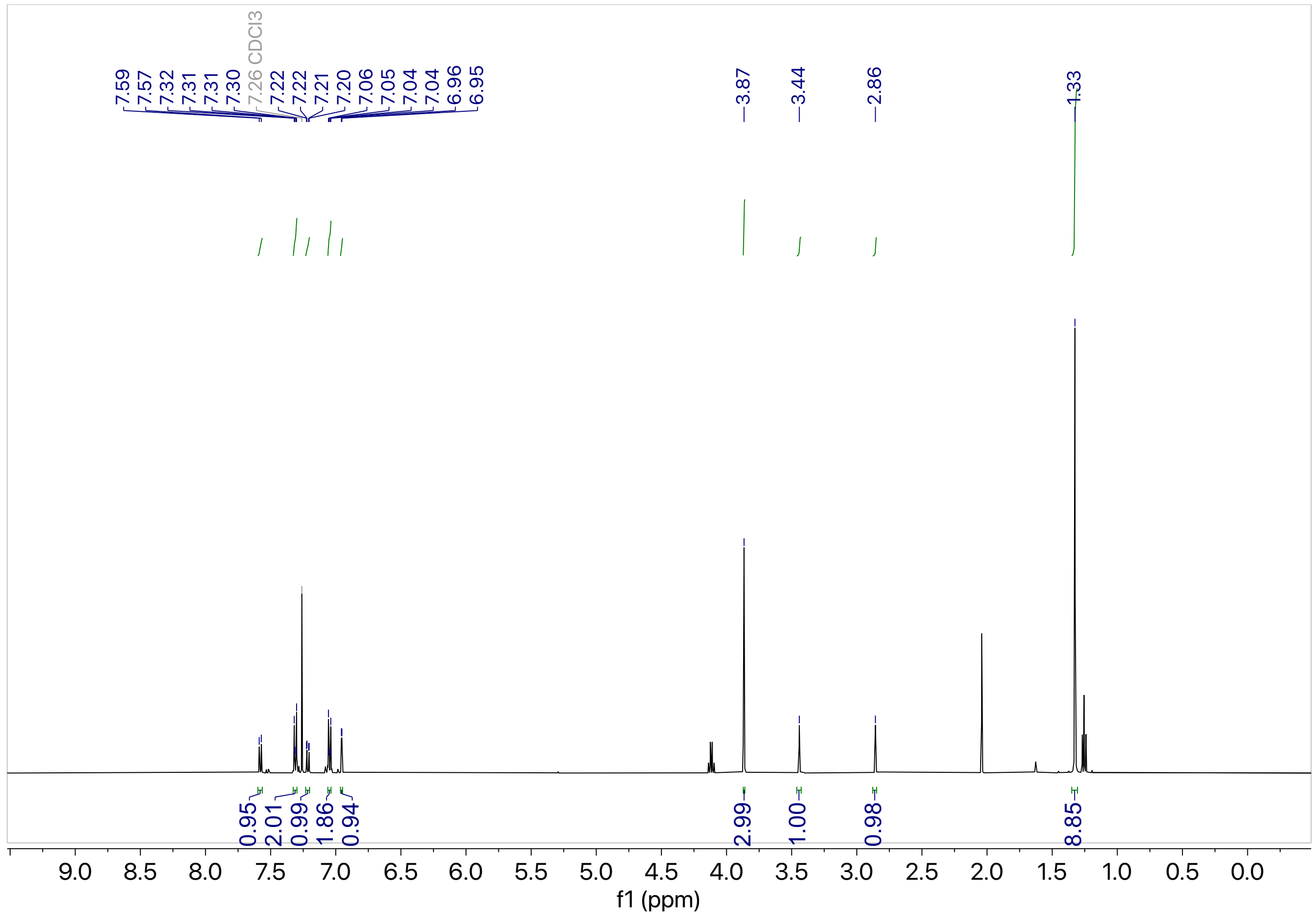

7.59
7.57
7.32
7.31
7.31
7.30
7.26 CDCl3
7.22
7.22
7.21
7.20
7.06
7.05
7.04
7.04
6.96
6.95
3.87
3.44
2.86
1.33
0.95
2.01
0.99
1.86
0.94
2.99
1.00
0.98
8.85
9.0
8.5
8.0
7.5
7.0
6.5
6.0
5.5
5.0
4.5
4.0
3.5
3.0
2.5
2.0
1.5
1.0
0.5
0.0
f1 (ppm)

**KL6-167B**

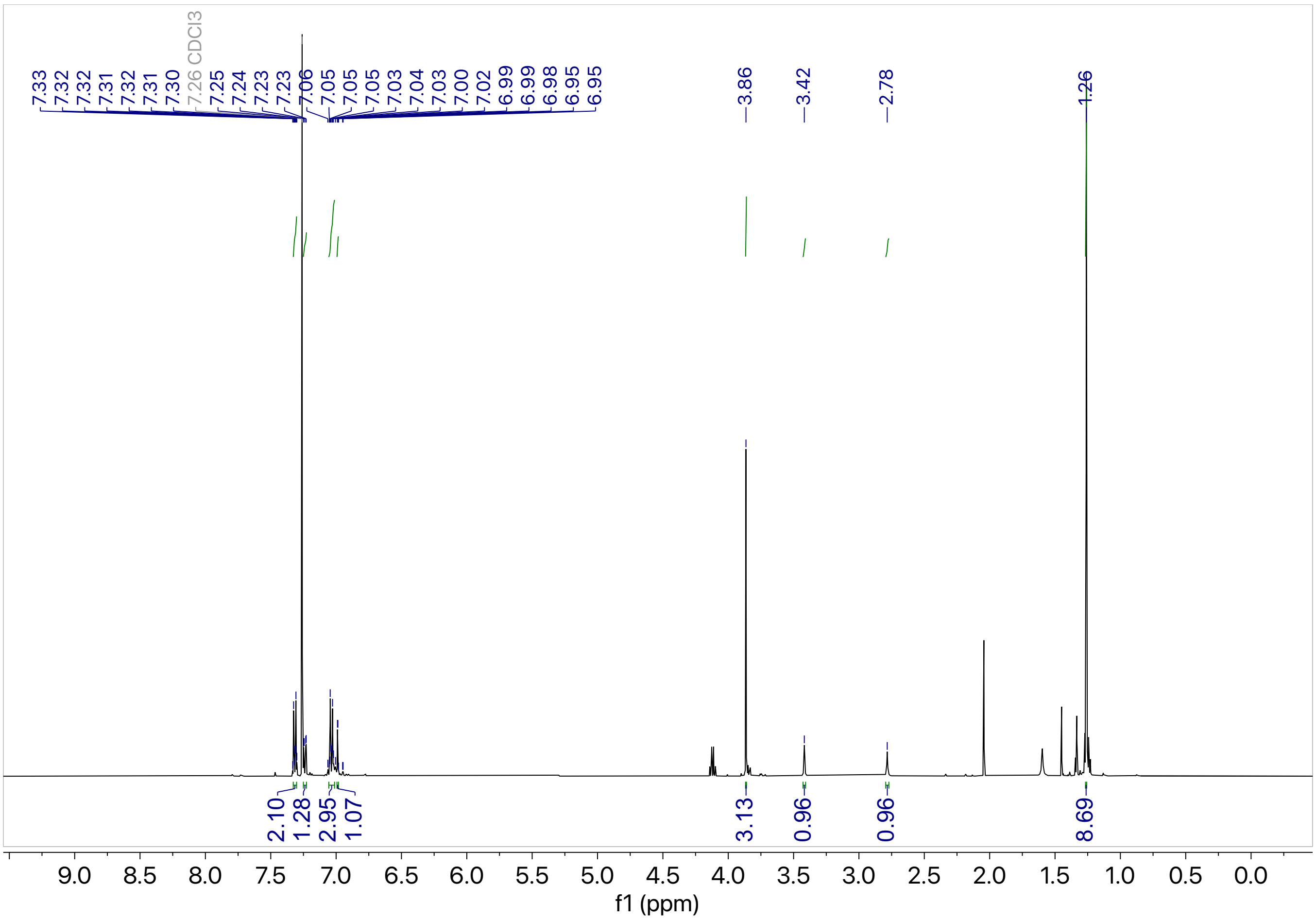

7.33
7.32
7.32
7.31
7.32
7.31
7.30
7.26 CDCl3
7.25
7.24
7.23
7.23
7.06
7.05
7.05
7.05
7.03
7.04
7.03
7.00
7.02
6.99
6.99
6.98
6.95
6.95
3.86
3.42
2.78
1.26
2.10
1.28
2.95
1.07
3.13
0.96
0.96
8.69
9.0 8.5 8.0 7.5 7.0 6.5 6.0 5.5 5.0 4.5 4.0 3.5 3.0 2.5 2.0 1.5 1.0 0.5 0.0
f1 (ppm)

**KL6-172A**

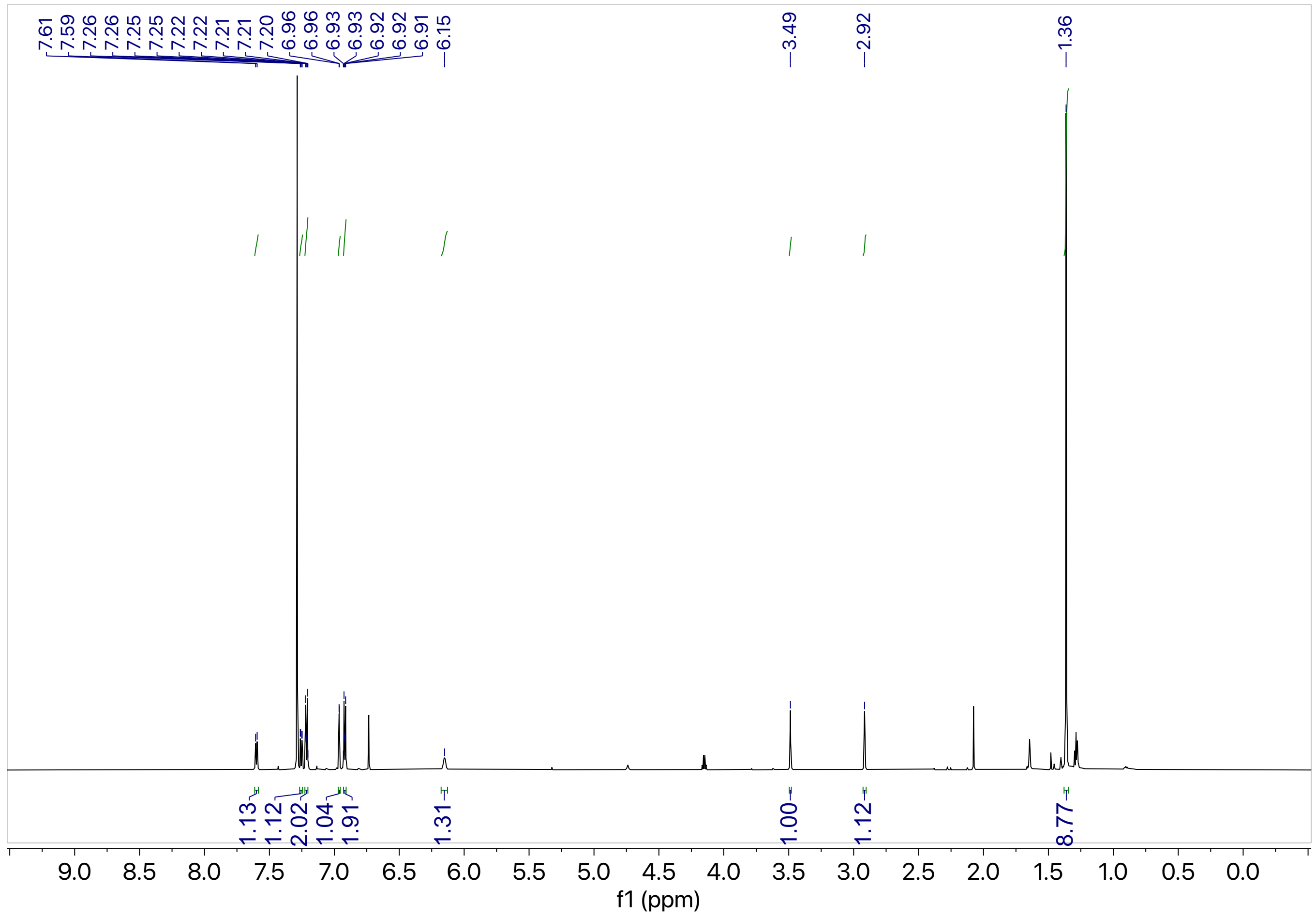

7.61
7.59
7.26
7.26
7.25
7.25
7.22
7.22
7.21
7.21
7.20
6.96
6.96
6.93
6.93
6.92
6.92
6.91
6.15
3.49
2.92
1.36
1.13
1.12
2.02
1.04
1.91
1.31
1.00
1.12
8.77
9.0
8.5
8.0
7.5
7.0
6.5
6.0
5.5
5.0
4.5
4.0
3.5
3.0
2.5
2.0
1.5
1.0
0.5
0.0
f1 (ppm)

**KL6-172B**

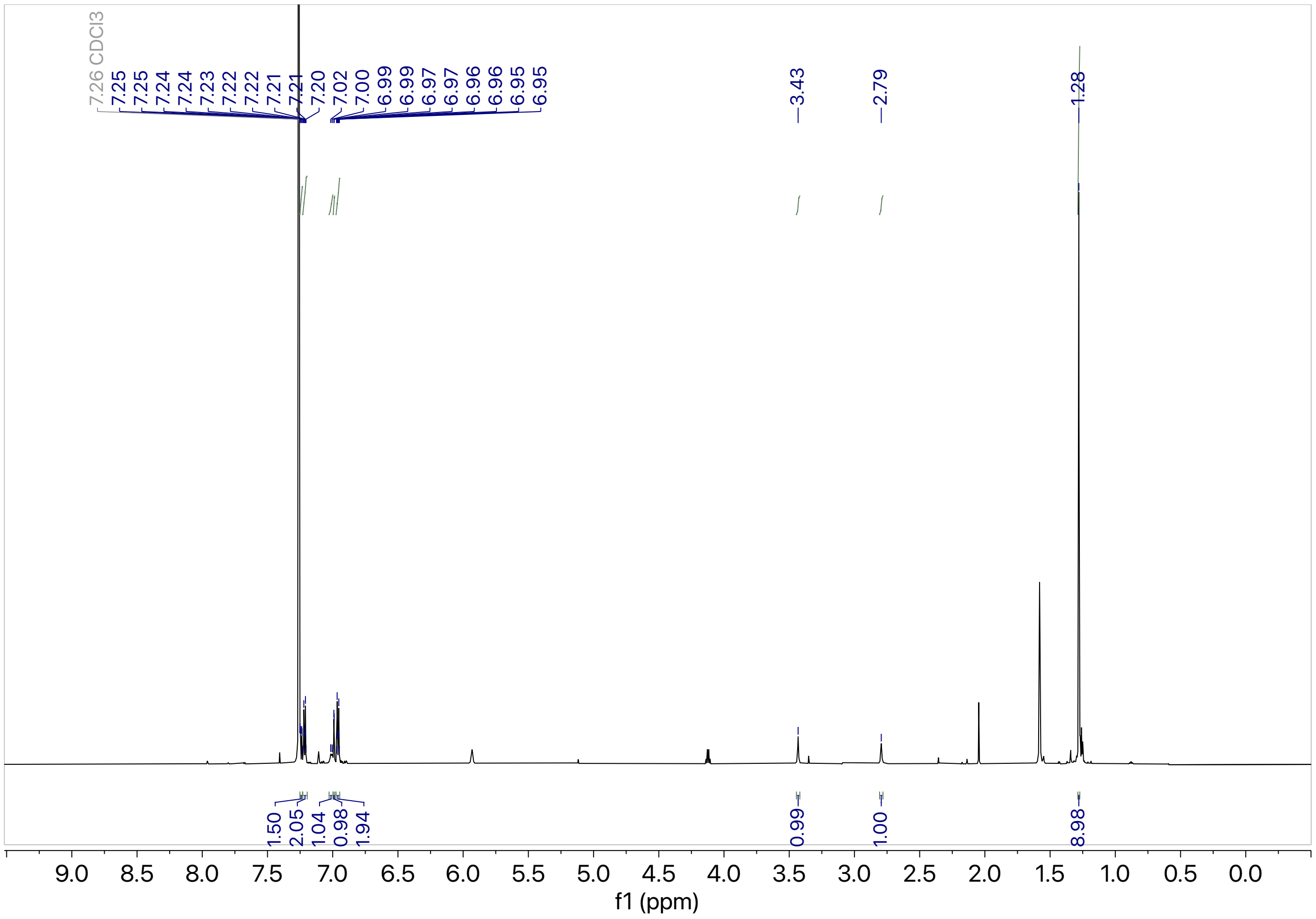
7.26 CDCl3
7.25
7.25
7.24
7.24
7.23
7.22
7.22
7.21
7.21
7.20
7.02
7.00
6.99
6.99
6.97
6.97
6.96
6.96
6.95
6.95
3.43
2.79
1.28
1.50
2.05
1.04
0.98
1.94
0.99
1.00
8.98
9.0
8.5
8.0
7.5
7.0
6.5
6.0
5.5
5.0
4.5
4.0
3.5
3.0
2.5
2.0
1.5
1.0
0.5
0.0
f1 (ppm)
